# An iPSC-derived bio-inspired scaffold modelling the structure and the effects of extracellular matrix in cardiac fibrosis

**DOI:** 10.1101/2024.02.07.578948

**Authors:** Francesco Niro, Soraia Fernandes, Marco Cassani, Monica Apostolico, Jorge Oliver-De La Cruz, Daniel Pereira- Sousa, Stefania Pagliari, Vladimir Vinarsky, Zbyněk Zdráhal, David Potesil, Vaclav Pustka, Giulio Pompilio, Elena Sommariva, Davide Rovina, Angela Serena Maione, Luca Bersanini, Malin Becker, Marco Rasponi, Giancarlo Forte

## Abstract

Cardiac fibrosis occurs following insults to the myocardium and is characterized by the abnormal accumulation of non-compliant extracellular matrix (ECM), which compromises cardiomyocyte contractile activity and eventually leads to heart failure. This phenomenon is driven by the differentiation of cardiac fibroblasts (cFbs) into myofibroblasts and results in changes in ECM biochemical, structural and mechanical properties. The lack of predictive *in vitro* models of heart fibrosis has so far hampered the search for innovative treatments. Here, we devised a single-step decellularization protocol to obtain and thoroughly characterize the biochemical and micro-mechanical properties of the ECM secreted by activated cFbs differentiated from human induced pluripotent stem cells (iPSCs). We activated iPSC-derived cFbs to the myofibroblast phenotype by tuning basic fibroblast growth factor (bFGF) and transforming growth factor beta 1 (TGF-β1) signalling and confirmed that activated cells acquired key features of myofibroblast phenotype, like SMAD2/3 nuclear shuttling, the formation of aligned alpha-smooth muscle actin (α−SMA)-rich stress fibres and increased focal adhesions (FAs) assembly. Next, we used Mass Spectrometry, nanoindentation, scanning electron and confocal microscopy to unveil the characteristic composition and the visco-elastic properties of the abundant, collagen-rich ECM deposited by cardiac myofibroblasts *in vitro*. Finally, we demonstrated that the fibrotic ECM activates mechanosensitive pathways in iPSC-derived cardiomyocytes, impacting on their shape, sarcomere alignment, phenotype, and calcium handling properties. We thus propose human bio-inspired decellularized matrices as animal-free, isogenic cardiomyocyte culture substrates recapitulating key pathophysiological changes occurring at the cellular level during cardiac fibrosis.

## Introduction

Heart fibrosis is a pathological condition that leads to the remodelling of cardiac chambers following the clinical manifestation of different heart pathologies, strongly contributing to the failure of the pump [^1^]. Among the main players in fibrosis onset and progression are cardiac fibroblasts (cFbs). cFbs orchestrate the maintenance of the structural framework of the tissue and sense specific molecular signals during cardiac injury that lead to their phenotype switch into activated cFbs, or myofibroblasts [^2, 3^]. Myofibroblasts are characterized by a reinforced cytoskeleton, supermature focal adhesions (FAs) and alpha-smooth muscle actin (α-SMA) stress fibres [^4^]. This contractile phenotype allows them to migrate to the site of injury, where they coordinate the deposition of a stiff, non-compliant extracellular matrix (ECM) [^5,6^]. The newly formed ECM protects the heart wall from rupture and is pivotal in preserving organ function by enriching the injury site with matricellular proteins and growth factors which promote reparative and regenerative signalling. These include fibronectin (FN), periostin (POSTN), and osteonectin (SPARC), among others, in a process which resembles the early phases of heart development [^7,8,9^]. Since adult cardiac tissue lacks endogenous stem cells with *in vivo* regenerative potential, the loss of terminally differentiated contractile cells cannot be replenished [^10, 11,12^]. Moreover, during the late stage of fibrosis, myofibroblasts undergo apoptosis, the ECM stiffens, and the abundant matrix infiltrates between the remaining functional cardiomyocytes (CMs), thus leading to arrhythmic events which compromise the heart function [^7,13^]. Different strategies have been devised to modulate cardiac fibroblasts activation and reduce the symptoms of cardiac fibrosis with contrasting results. Among them, angiotensin-converting enzyme (ACE) inhibitors, renin-angiotensin-aldosterone system (RAAS) inhibitors and transforming growth factor-beta (TGF-β) inhibitors are the most noteworthy [^14^]. In addition, innovative therapies targeting both cardiomyocyte regenerative ability and the excessive ECM deposition by cFbs are under preclinical evaluation [^15^]. Nonetheless, the development of effective therapies to treat cardiac fibrosis lags behind [^16,17,15^] in part due to the lack of *in vitro* models of myocardial fibrosis able to recapitulate human pathophysiology while capturing the patient genetic background [^18^].

Induced pluripotent stem cells (iPSCs) can be differentiated into cardiac cells, including CMs, cFbs, endothelial cells and tissue-specific macrophages, and are regarded as a viable alternative to the use of animal models and primary cell lines [^19, 20, 21, 22, 23^]. iPSCs require a basement membrane substitute to adhere, survive and differentiate in culture. While commonly used recombinant proteins such as fibronectin, laminin, vitronectin and collagens represent expensive options which provide iPSCs with some of the biological cues needed to grow and differentiate, more complex basement membrane derivatives like Matrigel ™ and Cultrex ™ gained momentum as coating solutions for iPSCs culture and to obtain cardiomyocytes from them. However, although proving necessary for iPSC propagation, they are generated from rodent tumours and thus lack the specific biological and mechanical cues of cardiac ECM [^24,25,23,26^]. In this context, organ decellularization represents an appealing opportunity to provide tissue-specific properties, but their use is hampered by the lack of donors and interspecies differences. More importantly, to the best of our knowledge, none of the options available on the market so far offers the possibility to generate isogenic models of human disease [^27, 28, 29,30, 31, 32, 33^].

To address the current challenges, we obtained human cardiac-specific decellularized matrices (dECMs) from cardiac fibroblasts (cFbs) differentiated from human induced pluripotent stem cells (iPSCs) and used them as culture substrates from cardiomyocytes generated from the same iPSC line. By modulating anti- and pro-fibrotic pathways, i.e. basic fibroblast growth factor (bFGF) and transforming growth factor beta 1 (TGF-β1) pathways, we demonstrated that it is possible to obtain fibrotic dECMs from activated iPSC-cFbs. After showing that the matrices are suitable for long-term culture of iPSC-derived cardiomyocytes (iPSC-CMs) [^34, 35, 36^], we generated an isogenic patient-specific model of fibrosis and unveiled characteristic changes of cardiac ECM composition, architecture and biomechanics induced by cFbs activation. Finally, we revealed how the iPSC-cFbs-derived fibrotic ECM affects iPSC-CMs morphology, phenotype, sarcomere assembly and calcium dynamics.

## Material and Methods

### iPSCs maintenance

The human iPSC cell line DF 19-9-7 T (iPSC, karyotype: 46, XY), purchased from WiCell (Madison, WI, USA), has been maintained in culture on Matrigel®-coated plates (1:100 in DMEM/F12, Corning) in Essential 8™ Medium (E8, Thermo Fisher Scientific) containing penicillin/streptomycin (P/S) (0.5%, VWR) and split 1:10 when they reached 80% of confluency. On the day of passage, E8 medium was supplemented with Y27632 (ROCK Inhibitor, RI) (10 μM, Selleck Chemicals, Houston, TX, USA) for the following 24 hours and then replaced with standard E8 media.

### Differentiation of iPSCs into CMs

iPSCs have been differentiated into CMs according the protocol from Lian et al., with slight modification [^19^]. In detail, 5×10^6^ iPSCs were detached by TrypLE™ Express Enzyme (1X), no phenol red (Gibco) and seeded on a Matrigel ®-coated 12-well plate in E8 supplemented with RI (10µM). The following day, RI was removed and the cells were grown in E8 for 2-3 days or until they reached 90% of confluence. Afterwards, the cells were cultured in RPMI 1640 with L-Glutamine (Biosera), P/S, Geltrex™ LDEV-Free Reduced Growth Factor Basement Membrane Matrix (1:100, Gibco), with the addition of B-27™ minus insulin (1×, Thermo Fisher Scientific) and supplemented with CHIR99021 (12 µM, Sigma-Aldrich), in order to modulate the Wnt/β-catenin signalling pathway. After twenty-four hours, CHIR activity was blocked by replenishing media with RPMI + B27 minus insulin and Geltrex (1:100). On day 3 of differentiation, a conditioned medium was prepared by combining one part of fresh RPMI B27 minus insulin with one part of medium taken from the cell culture plate and supplementing the obtained mix with Geltrex and IWP2 (5 µM, Selleck Chemicals). On day 5, the medium was replaced with RPMI + B27 minus insulin, and on day 7, the same medium was substituted with fresh RPMI supplemented with B-27™ (Thermo Fisher Scientific). Beating cells usually appeared between days 8 and 12 of differentiation, and were then used on day 20 for downstream experiments.

### Differentiation and maintenance of iPSC-cFbs

To differentiate iPSCs into cFbs, 5×10^6^ pluripotent cells were seeded on a Matrigel®-coated 6-well plate in E8 medium supplemented with RI (10 µM). The following day, medium was replaced with fresh E8 medium. At 80% confluency, the differentiation was started using a slightly modified version of the Zhang and colleagues protocol [^20^]. Briefly, on day 0 of differentiation, E8 medium was substituted with RPMI, B-27 minus insulin and CHIR99021 (12 µM). After 24 hours, CHIR99021 was removed by adding fresh RPMI + B27 minus insulin to the cells. On day 2, medium was further exchanged with FibroGRO™ Complete Media Kit for Culture of Human Fibroblasts (Merck), consisting of a fibroblasts-specific basal media supplemented with P/S (0.5%), L-glutamine (7.5 mM), ascorbic acid (AC) (50 µg/mL), hydrocortisone hemisuccinate (HC) (1 µg/mL), insulin (INS) (5 µg/mL), and HLL Supplement (HAS 500 µg/mL, linoleic acid 0.6 µM, and lecithin 0.6 µg/mL). Additionally, FibroGRO formulation was enriched with 75 ng/mL of Recombinant Human basic-FGF (bFGF, Peprotech). Medium was replaced every two days until day 20 of differentiation, when 1×10^6^ iPSC-cFbs were detached using TrypLe ™ and seeded onto 0.1% Gelatin (Sigma-Aldrich) T75 flask-culture coated plates (Corning) in FibroGRO™ Complete Media Kit supplemented with bFGF (75 ng/mL) and 2% of foetal bovine serum (FBS, Sigma-Aldrich). Only iPSC-cFbs with a passage number ≤ 10 (P10) were used for all the experiments.

### iPSC-cFbs culture on soft and stiff substrates

iPSC-cFbs were cultured onto T75 flask culture plates until they reached 80-90% of confluency. Next, cells were detached and seeded at a density of 2.5 × 10^5^ cells/cm^2^ either onto Corning® Costar® TC-Treated 24-well Plates (Corning) or 0.5 kPa and 64kPa CytoSoft® Imaging 24-well Plates (Advanced Biomatrix) in FibroGRO 2% FBS. Given the hydrophobicity of Polydimethylsiloxane (PDMS) substrates which characterize 0.5 and 64 kPa plates, to allow the correct cell adhesion, both CytoSoft and TCPS were coated with 5 µg/mL of Collagen Type I (SantaCruz). iPSC-cFbs were grown onto TCPS and CytoSoft plates for 7 days, changing media each second day, before downstream analysis.

### TGF-β1 treatment

To assess the effect of TGF-β1, iPSC-cFbs grown until 80-90% of confluency onto T75 flasks were detached and seeded on uncoated TC-treated multi-well plates at a cell density of 2.5 × 10^5^ cells/cm^2^. iPSC-cFbs were alternatively fed with FibroGRO 2% FBS alone, or supplemented with 10 ng/mL of TGF-β1 (Peprotech). Medium was changed each second day and cells cultured for 7 days, before downstream analysis.

### iPSC-cFbs activation assay

For the activation assay, iPSC-cFbs were detached from T75 flask and re-plated either onto Ibidi USA µ-Slide 8 Well, ibiTreat - Tissue Culture Treated Polymer Coverslip (Ibidi) or TC-treated multi-well plates without any coating material. iPSC-cFbs seeding concentration has been optimized according to the treatment used, given that cellular proliferation rate was affected by the administered drugs [^37, 38^]. iPSC-cFbs were seeded at a concentration of (i) 1.5×10^5^ cells/cm^2^ for ctrl condition, consisting in culturing the iPSC-cFbs in FibroGRO 2% FBS supplemented with 75 ng/mL bFGF, as during the maintenance; (ii) 2.5×10^5^ cells/cm^2^ for pro-fibrotic condition in which FibroGRO 2% FBS medium lacked bFGF supplementation; and (iii) 1.0×10^5^ cells/cm^2^ for anti-fibrotic condition consisting in FibroGRO 2% FBS supplemented with 75ng/mL of bFGF and 500nM of TGF-β inhibitor (A83-01, Tocris). iPSC-cFbs were maintained for 7 days in culture replacing medium each second day. Downstream analyses were performed by using 100% confluent cells.

### Decellularization procedure and storage of dECM

To isolate the ECM produced by cFbs, confluent iPSC-cFbs were washed in PBS and incubated for 5 minutes at 37 °C in PBS containing NH4OH (20 mM, Sigma-Aldrich) and 0.1% Triton X-100 (Sigma-Aldrich) to induce cell degradation. The dECM produced by iPSC-cFbs (cFbs-dECM) were washed three times with PBS and incubated for 30 minutes at 37 °C with 60 ng/mL DNAse I (Stemcell technologies) in DNAse buffer solution (Table 1). Next, dECM were washed five times with PBS and either used immediately or stored at 4 °C for future use (up to 5 days).

**Table 1.**
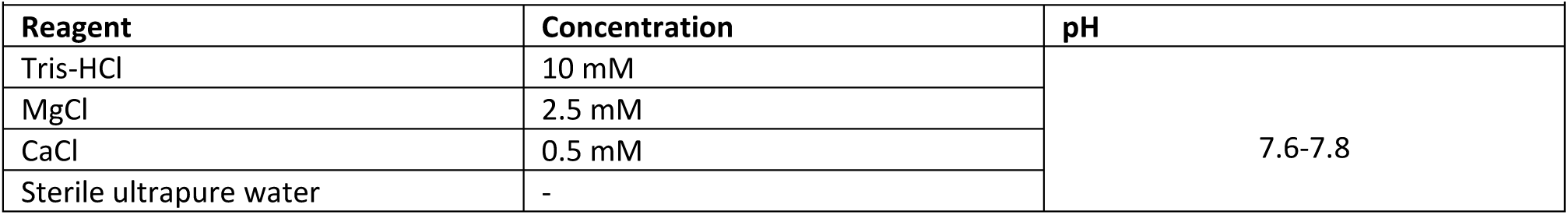
–DNAse buffer composition.

### iPSC-CMs seeding and culturing on dECM

iPSC-CMs on day 20 of differentiation were detached by TrypLe ™ from Matrigel ®-coated 12-well plates and seeded for single-cell experiments at a cell density of 2.0×10^5^ cells/cm^2^ onto µ-Slide 8 well ibiTreat containing adherent cFbs-dECM in RPMI supplemented with B27. Alternatively, iPSC-CMs were split 1:1 onto cFbs-dECM coated TC-treated 12-well plates. iPSC-CMs were grown onto cFbs-dECM for 10 days, changing media each third day, before downstream analysis.

### RNA extraction, Reverse Transcription, and Real Time quantitative PCR analysis

RNA extraction from cultured cells has been performed using the High Pure RNA Isolation Kit (Roche) according to the manufacturer’s protocol. RNA has been retrotranscribed into complementary DNA (cDNA) using the RT2 674 First 675 Strand Kit (SA Biosciences, Frederick, USA). Gene expression levels were quantified by Reverse Transcription quantitative polymerase chain reaction (RT–qPCR) using the LightCycler 480 Real-Time PCR System (Roche, Basel, Switzerland). Cycling parameters were set at (i) 95 °C for 10 minutes (1 cycle), and (ii) 95 °C for 15 seconds followed by 60 °C for 1 minute (45 cycles). Results were normalized on the expression of the housekeeping gene glyceraldehyde-3-phosphate dehydrogenase (GAPDH), and expressed as log102^-ΔΔCt^. Primers used are listed on Table2.

### Flow Cytometry

Cells were detached and washed once in FACS buffer (UltraPure™ 0.50mM EDTA -Gibco-with 0.5% FBS in PBS), and fixed with IC Fixation buffer (eBioscience™) for 20 minutes at room temperature (RT) in the dark. Next, cells were washed first in FACS buffer and subsequently in Permeabilization Buffer 10X (eBioscience™) diluted 1:10 (1X) in ultrapure sterile water. Cells were then blocked in 2.5% bovine serum albumin (BSA, Sigma-Aldrich) diluted in 1X permeabilization buffer. Next, cells were incubated with primary antibody (or conjugated antibody) diluted in 1% BSA - 1X permeabilization buffer for 30 minutes in the dark at RT. Cells were then washed with 1X permeabilization buffer and incubated again for 30 minutes with a secondary antibody at RT in the dark. After the incubation, cells were washed twice, once with 1X permeabilization buffer and once with FACS buffer, and finally resuspended in 200µL of FACS buffer before being analysed on a BD FACS Canto™ (BD Biosciences). The samples were analysed using the FACSAria II (Becton-Dickinson, USA), and plots were prepared with FlowJo software V10 (Tree Star, USA).

### Immunocytochemistry

To perform immunofluorescence, the samples were washed three times in PBS, fixed in 4% paraformaldehyde (PFA) solution for 12 minutes at room temperature and rinsed three times in PBS. Cells were then permeabilized using 0.1% Triton X, diluted in PBS, for 10 minutes. Following permeabilization, cells were washed three times in PBS and blocked in 2.5% BSA for 30 minutes at room temperature. Cells were incubated with primary antibody, diluted in 1% BSA, overnight at 4 °C, or alternatively for 2 hours at RT (Table 2). Afterwards, cells were washed three times and incubated for 1 hour in Alexa fluorochrome-conjugated (AF) secondary antibodies 488/555/647 diluted in PBS (1:500, Thermo Fisher Scientific) at RT. F-actin was stained with AF-647 conjugated phalloidin diluted in PBS (1:500, Thermo Fisher Scientific) for 1 hour at RT. At the end of incubation, cells were washed three times and counterstained with 4‟,6‟-diamidino-2-phenylindole (DAPI, Sigma-Aldrich) for 5 minutes at RT. Next, cells were washed three times with PBS, and embedded in Moviol mounting medium (Merck). Images have been acquired using a Zeiss LSM 780 confocal microscope at an optimal interval suggested by the software, and visualized by applying the maximum intensity projection algorithm. Detailed pictures of cell structures have been captured using super resolution microscopy (Elyra 7, Zeiss) with structured illumination microscopy (SIM) mode (CEITEC, CELLIM Facility).

**Table 2.**
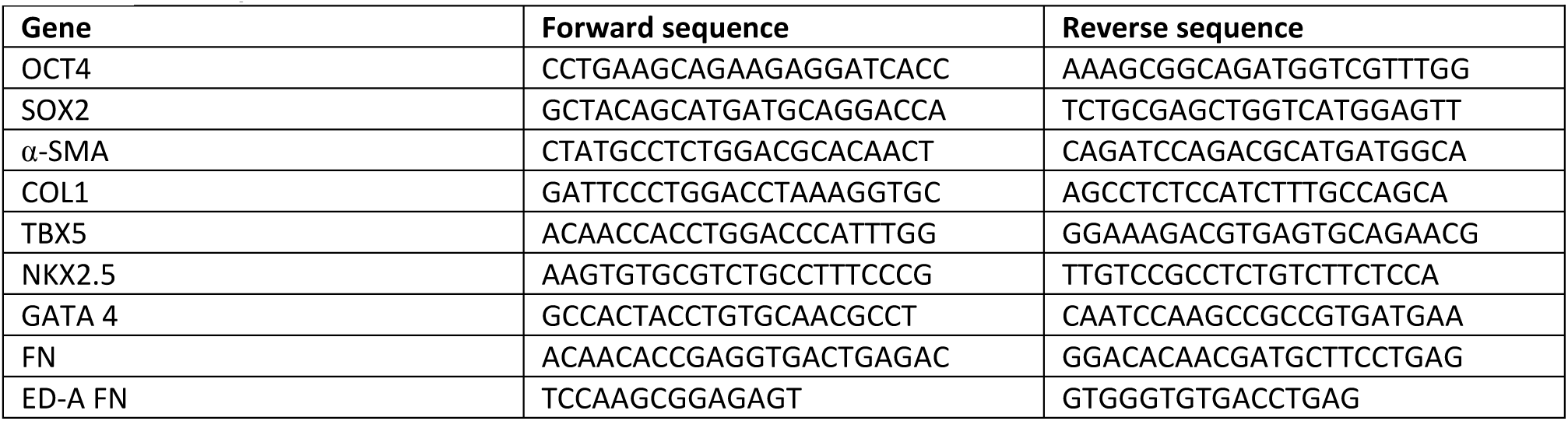
– List of primers.

### Image analysis

The images acquired through Zeiss confocal microscopy were processed as described in the following subsections.

_ The percentage of GATA4 and SMAD2/3 positive cells was identified by a customized pipeline run on Imagej-win64 and Cell Profiler Analist ™ 4.2.6 to automatically count cell nuclei and nuclear transcription factors in iPSC-cFbs [^39^].

_ iPSC-cFbs stress fibres alignment was quantified by the ImageJ-win64 plugin, OrientationJ, which allowed the identification of α-SMA fibres into a dominant direction (coherency – 0-1 arbitrary unit) [^40, 41^].

_ Cell area has been manually calculated marking the cellular borders of iPSC-cFbs on Imagej-win64.

_ The number of FAs and their area were calculated on iPSC-cFbs (marked for vinculin), using a modified version of the algorithm described by Horzum and colleagues [^42^].

_ CMs morphological parameters (cellular and nuclear area, and circularity), as well as their protein localization (nucleus-cytoplasm ratio) were calculated applying a customized pipeline on Imagej-win64 and Cell Profiler Analist ™ 4.2.6 [^43, 44, 39^]. Briefly, CMs were identified through the detection of DAPI and α-sarcomeric actinin (α-actinin) (AF-488) by which cellular and nuclei area, as well as circularity were calculated. Transcription factors localization was analysed quantifing the colocalization of transcription factors either with DAPI or α-actinin [^43, 44, 39^].

_ For sarcomere parameters analysis, iPSC-CMs cultured onto ctrl and pro-fibrotic dECM have been fixed and stained for α-actinin. Cells were imaged at high magnification (> 60x) in order to obtain detailed images of myocyte sarcomeres using Lightning mode in LEICA (TCSSP8) confocal microscope. Acquired images have been processed using the Imagej plugin for the automatic quantification of sarcomere assembling, SotaTool [^45^].

### Calcium handling measurements

Fluo-4 Calcium Imaging Kit (Thermo Fisher Scientific) was used for the detection of calcium spikes. Briefly, iPSC-CMs were incubated with 1 µM of Fluo-4 dye (ex/em 494/506 nm) in cell culture medium for 30 minutes at 37 °C. Medium was replaced with fresh pre-warmed medium, and the cells placed in an incubation chamber (37 °C and 5% CO2) under a Zeiss LSM 780 confocal microscope for 15 minutes to allow cell adaptation following media change. Beating clusters were selected, and calcium spikes were registered. The registrations of beating CM clusters were analysed by a customized pipeline involving Imagej and RStudio (https://posit.co/download/rstudio-desktop/), which allowed the detection of Fluo-4 intensity and spiking frequency. The output of the analysis revealed the parameter of interests relative to calcium handling (beating rate, raising - rT10-rT90 - and decay - dt- time).

### Enzyme linked immunosorbent assay

Cell culture medium was collected from iPSC-cFbs on day 6 of activation assay and centrifuged at 13000 rpm for 10 minutes. The supernatant was separated from the pellet and stored at -80 °C. The amount of TGF-β1 present in the media was quantified using the Human/Mouse/Rat/Porcine/Canine TGF-beta 1 Quantikine ELISA kit(R&D systems). The results were normalized to the total number of cells in the culture at the moment of media collection.

### Scanning electron microscopy of dECM

dECMs were fixed in 3% glutaraldehyde/100 mM cacodylate buffer (Sigma Aldrich) for 30 minutes at RT. Next, samples were washed 3 times in cacodylate buffer (10 minutes/wash), and dehydrated through washes in serial dilution of ethanol (10% to 100%). The dehydrated samples were coated for 15 minutes with a JFC-1300 autofine coater (JEOL) at 40 mA. Images have been acquired in high-vacuum conditions using a JCM-6000 benchwork SEM (JEOL). SEM was equipped with a SEI detector PC-standard, with accelerating voltage set at 15kV. Magnification was set at x2000.

### Mass-spectrometry: sample preparation and analysis

Following decellularization, dECM samples were prepared for mass spectrometry. In detail, after DNAse treatment dECMs from the three different conditions were incubated overnight at 4 °C with 5% acetic acid. To extract the proteins, the dECMs were harvested using a cell scraper and collected into a tube containing the Extraction buffer described in Table 3. The extracted samples were stored at -80 °C and prepared for Liquid Chromatography with tandem mass spectrometry (LC-MS/MS) by filter-aided sample preparation (FASP) method [^46^]. The final peptide mixture (500 ng) was separated by a 104-minute long linear LC gradient (Ultimate 3000 RSLCnano, Thermo Fischer Scientific) and analysed by timsTOF Pro mass spectrometer (Bruker). Data were acquired in data-independent acquisition (DIA) mode in the precursor m/z range of 400-1000. DIA LC-MS data were processed using the DIA-NN application (version 1.8; https://github.com/vdemichev/DiaNN). The protein library has been generated following cRAP contaminant (http://www.thegpm.org/crap/) and UniProtKB-Human databases (ftp://ftp.uniprot.org/pub/databases/uniprot/current_release/knowledgebase/reference_proteomes/Eukaryota/UP0 00005640/UP000005640_9606.fasta.gz), and match between runs (MBR) across the whole dataset. Database search results were set to follow the false discovery rates (FDR) thresholds (precursor level and protein group level: 1% FDR). The default protein inference algorithm implemented in DIA-NN was used to construct the protein groups list utilizing proteotypic (i.e. peptides unique for the given protein within the whole protein database) and non-proteotypic peptides. Data were normalized on the ctrl dECM using KNIME software (https://www.knime.com/) applying the MaxLFQ algorithm and represented as log2 fold change (FC) of protein intensities. Statistics were calculated with a moderated t-test using the LIMMA package in R (https://bioconductor.org/packages/release/bioc/html/limma.html for more details), and p values adjustment was done using the Benjamini & Hochberg method. For the analysis, R software version 4.3.0 and the biplot function in the PCATools package version 2.12.0 were used to generate the principal cluster analysis (PCA) plot containing the dECM proteins contributing to the main differences between ctrl, pro-, and anti-fibrotic dECM. The heatmap was generated using Heatmapper software (http://www.heatmapper.ca/) by uploading PG intensities of the different dECM components, and they are represented as Row Z-score. Biojupies web tool (https://maayanlab.cloud/biojupies/analyse) has generated the standard PCA starting from the differentially expressed matrisome proteins. The volcano plots were generated using Bioconductor software (https://bioconductor.org/packages/release/bioc/html/limma.html). Bar plots representing gene ontology studies were obtained from the EnrichR database (https://maayanlab.cloud/Enrichr/) by uploading the 37 significantly expressed matrisome proteins between ctrl and pro-fibrotic dECM obtained following moderated t-test using LIMMA. The mass spectrometry proteomics data have been deposited to the ProteomeXchange Consortium via the PRIDE [^47^] partner repository with the dataset identifier PXD049208.

**Table 3.** – List of primary antibodies.

| Table 3 – List of primary antibodies |  |  |  |  |
| --- | --- | --- | --- | --- |
| Method | Target protein | Catalog number | Dilution | Supplier |
| Immunofluorescence | PDGFR-α | sc-398206 | 1:100 | Santa Cruz |
|  | Fibroblast-specific-protein | F4771 | 1:200 | Sigma-Aldrich. |
|  | DDR2 | ab63337 | 1:200 | Abcam |
|  | TE-7 | CBL-271 | 1:100 | Merck |
|  | GATA4 | 36966 | 1:400 | Cell Signaling |
|  | Alpha-smooth muscle actin | ab7817 | 1:200 | Abcam |
|  | F-actin | A22287 | 1:200 | ThermoFisher SCIENTIFIC |
|  | Vinculin | V9131 | 1:200 | Sigma-Aldrich |
|  | SMAD 2/3 | 8685 | 1:400 | Cell-Signalling |
|  | COL 1 | 600-401-103 | 1:100 | Rockland Immunochemicals |
|  | COL 4 | PA5-104508 | 1:50 | ThermoFisher SCIENTIFIC |
|  | Fibronectin | 26836 | 1:800 | Cell Signaling |
|  | Extra-domain A fibronectin | sc-5986 | 1:100 | Santa Cruz |
|  | Extra-domain B fibronectin | ab154210 | 1:100 | Abcam |
|  | Laminin | L9393 | 1:200 | Sigma-Aldrich |
|  | Alpha-sarcomeric actinin | A7811 | 1:300 | Sigma-Aldrich |
|  | YAP | 14074 | 1:200 | Cell Signalling |
|  | MRTF-A | 14760 | 1:200 | Cell Signaling |
|  | NFAT1 | 4389 | 1:200 | Cell Signaling |
|  | Ki67 | ab16667 | 1:500 | Abcam |
| Flow cytometry | CD31-FITC | 11-0311-82 | 1:20 | ThermoFisher SCIENTIFIC |
|  | CD90-APC | 1A-625-T100 | 1:10 | Exbio |
|  | CD45-APC | 17-0451082 | 1:10 | ThermoFisher SCIENTIFIC |
|  | CD73-FITC | 41-0200 | 1:10 | ThermoFisher SCIENTIFIC |
| | α-SMA-FITC | ab7817 | 1:10 | Abcam |
|  | FAP-APC | FAB3715A | 1:10 | RnD Systems |
|  | Cardiac TNNT-FITC | 130-119-575 | 1:50 | Miltenyi Biotec |
|  | Ki67 | ab156956 | 1:10000 | Abcam |

**Table 4.** – Extraction buffer composition.

| **Reagent** | **Concentration** | **Supplier** |
| --- | --- | --- |
| Tris-HCL pH 6,8 | 125 mM | ThermoFisher SCIENTIFIC |
| Sodium dodecyl sulfate (SDS) | 1% | Sigma-Aldrich |
| DL-dithiothreitol (DTT) | 20 mM | ThermoFisher SCIENTIFIC |
| Glycerol | 10% | Sigma-Aldrich |
| Phenylmethanesulfonyl fluoride (PMSF) | 0.05 mM | Sigma-Aldrich |
| Protease inhibitor | 1x | Santa Cruz |
| EDTA | 1 mM | Sigma-Aldrich |
| Sterile ultrapure water |  | - |

### dECM indentation and topographic/mechanical parameters measurements

Nanoindentation measurements were performed with an interferometer-based indentation device (Pavone, Optics11 life). All measurements were performed at RT with samples fully hydrated by immersion in PBS. Samples were analysed using a probe with a 9.5 µm tip radius and stiffness of 0.49 N/m and a tip with a 24 µm radius and stiffness of 0.018 N/m, dependent on sample stiffness and topography. Samples with inhomogeneities in surface coverage of dECM (assessed by built-in brightfield microscopy) were indented on spots where the presence of ECM was visually confirmed to avoid probing the bare well plate or skinny layer ECM, as this would not fulfil the fitting requirements. For each condition, Young’s modulus of n ≥ 30 individual spots from N ≥ 2 wells was investigated by fitting the indentation curves between 0 nm and 500 nm indentation depth, following the Hertzian model with the software provided by the Pavone manufacturer and assuming a Poisson ratio of 0.5. Fits with an R2 < 0.8 were excluded. Based on these fits, the surface topography could be assessed by comparing the relative z-position determined as the contact point for matrix scans, where a z=0 µm is assumed to represent the well plate surface. Dynamic measurements were performed at n ≥ 3 spots with clearly visible matrix at 1000 nm indentation depth with an amplitude of 100 nm at 1 Hz, 10 Hz, and 20 Hz to investigate the frequency-dependent visco-elastic behaviour.

### Statistical analysis

Statistical analysis was performed on at least three biological (N) or technical (n) replicates. Results were analysed with GraphPad Prism v. 6.0 (San Diego, USA). Specific statistical analyses have been performed depending on the conditions used in the experiments, and they are indicated in figure legends. A p < 0.05 was considered statistically significant.

## Results

### Generation of iPSC-cFbs with a pro-fibrotic phenotype

cFbs are the major contributors of heart ECM physiological homeostasis and pathological remodelling [^2, 3^]. We exploited a protocol to derive cells displaying the morphological and the phenotypical signature of cFbs [^20^]. Briefly, pluripotent cells were stimulated to commit to cardiac mesoderm by regulating the Wnt signalling pathway through the glycogen synthase kinase 3 (GSK3α) inhibitor, CHIR-99021, and then prompted with bFGF to acquire a cardiac fibroblast-like phenotype (Figure 1A). After 20 days of culture, the differentiated iPSC-cFbs were characterized and used for further experiments. Gene expression analysis performed by RT-qPCR demonstrated that the differentiation protocol consistently yielded the down-regulation of pluripotency genes (OCT4, SOX2) and the up-regulation of fibroblast (α-SMA and collagen 1a1, COL1A1) and cardiogenic (GATA4, TBX5, NKX2.5) markers [^48, 49, 50^] (Figure 1B). We confirmed these results by performing immunohistochemistry with antibodies directed against well-established fibroblast markers, such as fibroblast specific protein (FSP), TE-7, platelet-derived growth factor receptor-α (PDGFR-α) and discoidin domain receptor 2 (DDR2, Figure 1C). This analysis confirmed that the cells generated by the differentiation protocol displayed a spindle-shaped morphology, a well-known feature of fibroblasts. Noteworthy, the same analysis revealed that 90.06 ± 0.46% of cells derived were positive for the cardiac transcription factor GATA4 (Figure 1C and 1D) [^51^]. The purity of the generated cells was assessed by flow cytometry for the presence of the fibroblast marker CD90 (Figure 1E). The results showed that 84.14 ± 9.47 % of the cells were positive for CD90 (Figure 1F), while negligible levels of endothelial (CD31) and hematopoietic cell markers (CD45) were detected (3.71 ± 3.91% and 1.81 ± 2.08% respectively, Figure 1E and 1F). Variable levels of mesenchymal progenitor marker CD73 (21.90 ± 37.12%) were also found (Figure 1E and 1F) [^52^]. Altogether, these results confirmed the efficiency of the protocol used for cFbs differentiation.

**Figure 1.**
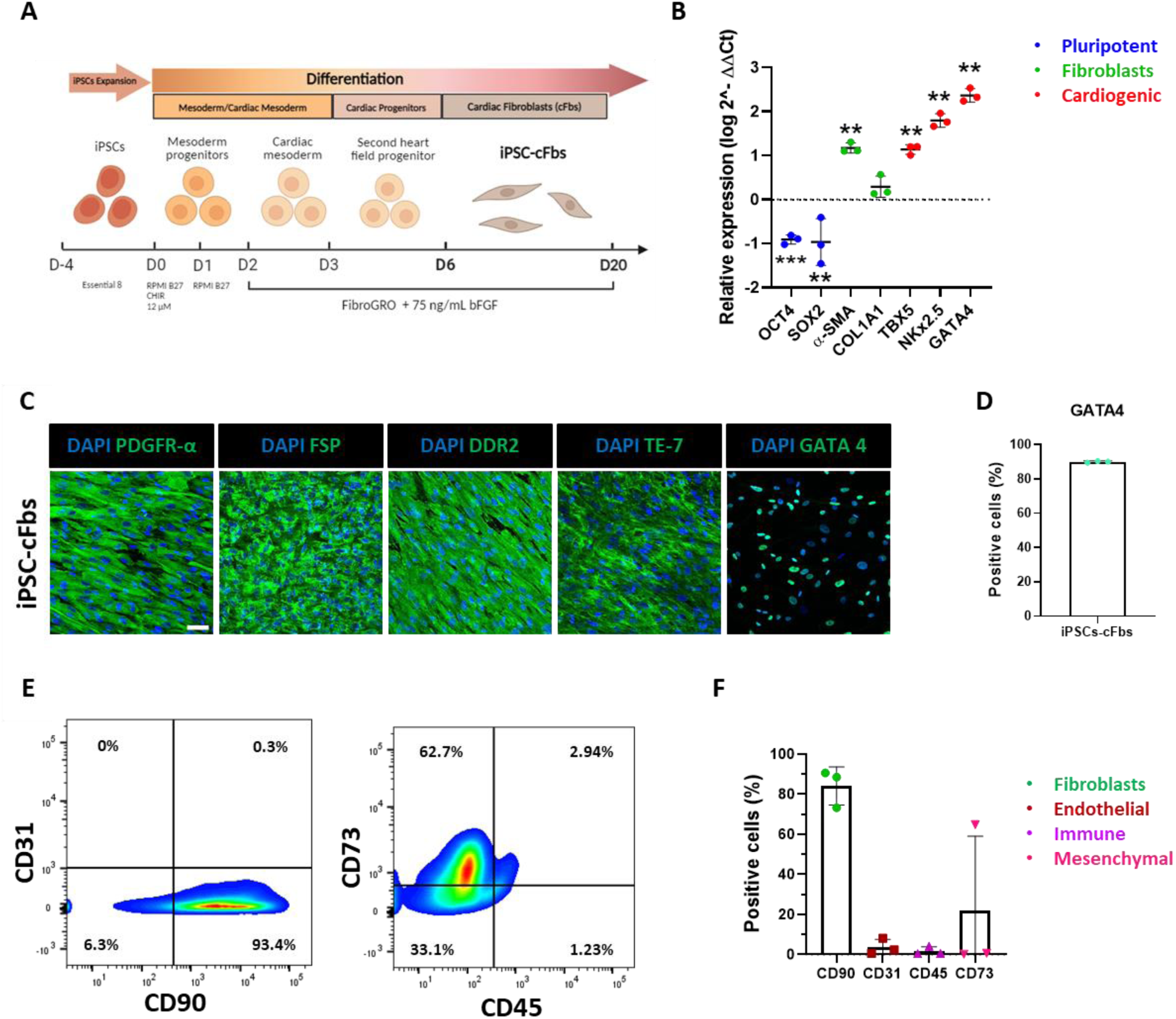
Cardiac fibroblasts derived from express fibroblast- and cardiac-specific markers. **(A)** Schematic overview of the differentiation protocol used to obtain cardiac fibroblasts from induced pluripotent stem cells. **(B)** Gene expression analysis of selected pluripotency (blue), fibroblast (green), and cardiac (red) genes in iPSC-derived cardiac fibroblasts as obtained from RT-qPCR. Gene expression values are expressed as fold change (log 2^-ΔΔCt, FC) compared to iPSCs and normalized to iPSCs. Statistical analysis was performed with One-sample t-test (N=3); **p<0.01, ***p < 0.001. Data presented as mean ± standard deviation. **(C)** Representative confocal images of iPSC-derived cardiac fibroblasts stained with antibodies directed against the indicated fibroblast-specific (PDGFR-α, FSP, DDR2, TE-7) and cardiac (GATA4) markers (N=3). The stainings are shown in green and nuclei are counterstained with DAPI (blue) (Scale bar: 50µm). **(D)** Barplot representation of the percentage of GATA4-positive iPSC-derived cardiac fibroblasts (N=3) obtained from confocal imaging. Data presented as mean ± standard deviation. **(E)** Representative contour plot images of the expression of the indicated surface markers in iPSC-derived cardiac fibroblasts. **(F).** Barplot representation of the expression of the indicated markers as obtained from flow cytometry analysis. The values are expressed as percentage of positive cells (N=3). Data presented as mean ± standard deviation.

The activation of fibroblasts into myofibroblasts is triggered by the local availability of pro-fibrotic molecules like TGF-β1 and favoured by the stiffening of the surrounding microenvironment [^53,54^]. Previous reports demonstrated that primary cFbs become spontaneously activated when cultured *in vitro* on substrates with supraphysiological stiffness, like tissue culture polystyrene surfaces (TCPS, Young’s modulus > 1 GPa) [^55^]. We cultured iPSC-cFbs on substrates with increasing stiffness (0.5 kPa, 64kPa and TCPS) for 7 days and quantified the percentage of cells co-expressing α−SMA and fibroblast activated protein (FAP) as proxies for myofibroblast differentiation [^56,57^]. This experiment clarified that a significantly higher percentage of iPSC-cFbs co-expressed α−SMA and FAP when cultured on TCPS as compared to 0.5 and 64 kPa (Supplementary Figure 1A). Confocal microscopy demonstrated that cells cultured onto TCPS grew markedly bigger than those cultured onto 0.5 and 64 kPa substrates (Supplementary Figure 1B) and displayed the formation of aligned α−SMA-rich stress fibres typical of myofibroblasts (Supplementary Figure 1C). We also designed a control experiment where iPSC-cFbs were cultured on TCPS for 7 days in the presence of TGF-β1, the protein recognized as the main inducer of the myofibroblast contractile phenotype *in vivo [*^58^]. The results showed that αSMA expression in TGF-β-stimulated cells was not different from that found in the same cells grown on TCPS alone (Supplementary Figure 1D). RT-qPCR confirmed that the gene expression of α−SMA did not differ when compared with TCPS cells. Similar results were obtained when we used primers directed against COL1A1 gene and extra-domain A fibronectin splicing isoform (ED-A FN) (Supplementary Figure 1E). ED-A FN isoform expression represents one of the main markers of cardiac fibrosis involved in the reinforcement of FN multimeric complexes and in the activation of latent TGF-β [^59, 60, 61^]. These results indicated that TCPS stiff substrate *per se* induced the spontaneous activation of iPSC-derived cells, similar to what has been previously described in primary cFbs [^62^]. Furthermore, the activation was not affected by the addition of exogenous TGF-β1 stimulation.

Next, we set at identifying culture conditions suited to modulate iPSC-cFbs spontaneous activation on stiff TCPS substrate. bFGF is known to inhibit the activation of cFbs into myofibroblasts [^63,35^]. We thus repeated the experiment using TCPS to culture the iPSC-cFbs in the presence or absence of bFGF (75 ng/mL), alone or in combination with TGF-β receptor pharmacological inhibitor (A83-01, 500 nM) [^64^]. After 7 days, we analysed the percentage of cells co-expressing α-SMA and FAP by flow cytometry. This analysis confirmed that bFGF exclusion from media of cFbs growing on TCPS enhanced the co-expression of the two fibrotic proteins α-SMA/FAP (69.82 ± 8.11 % vs 46.47 ± 4.20 % in presence of bFGF). On the contrary, the combination of bFGF with A83-01 resulted in a significant reduction of α-SMA/FAP positivity to 19.77 ± 7.52 % (Figure 2A, B). Next, we used RT-qPCR to quantify the expression of genes characteristic of the activated phenotype. We confirmed that TCPS prompted a significant increase in the expression of α-SMA, COL1A1, FN and the ED-A FN genes as compared to the treatment with bFGF alone or in combination with A83-01 (Figure 2C). Altogether these results indicated that TCPS stiffness might act as a powerful pro-fibrotic stimulus for iPSC-cFbs, and that bFGF and the combination of bFGF and TGF-β receptor inhibitor A83-01 could be used to modulate their fibrotic phenotype. Hence, we named iPSC-cFbs cultured onto TCPS as pro-fibrotic, while those cultured in the presence of bFGF and A83-01 combination were defined as anti-fibrotic. Cells grown in the presence of bFGF alone were defined as control (ctrl) (Supplementary Figure 1F). Next, we imaged the iPSC-cFbs exposed to the various treatments using the super-resolution microscope. Image analysis revealed that the cells treated in pro-fibrotic culture conditions were significantly bigger than the control and the anti-fibrotic counterparts (Figure 2E). Activated myofibroblasts are known to assemble robust FA complexes that reinforce their ability to interact and remodel the surrounding ECM [^65^]. We thus stained FA protein vinculin (shown in Figure 2D) and quantified its expression and focal adhesion area in pro-fibrotic, ctrl and anti-fibrotic iPSC-cFbs (Figure 2F). The outcome confirmed that pro-fibrotic conditions induced the assembly of a significantly higher number of FAs in iPSC-cFbs as compared to the same cells cultured in anti-fibrotic or ctrl conditions (Figure 2F). Consistently, the size of the FAs was significantly higher in those cells compared to the ctrl or anti-fibrotic counterparts (Figure 2F). Our hypothesis was further strengthened by staining the cells for α-SMA (Figure 2G), indicating that the pro-fibrotic cells appeared well aligned and displayed oriented α-SMA-rich stress fibres which are characteristic of activated fibroblasts (Figure 2H). Finally, we quantified TGF-β1 release in the supernatant of iPSC-cFbs *via* ELISA and confirmed that cells grown in the presence of bFGF display reduced ability to release endogenous TGF-β1 (Figure 2I). As a consequence, the nuclear translocation of TGF-β1 downstream effector SMAD 2/3 (Figure 2J) [^66^] was significantly reduced in iPSC-cFbs cultured in the presence of ctrl or anti-fibrotic conditions (pro-fibrotic: 65.49±17.88%; ctrl: 17.13±7.25%; anti-fibrotic: 18.19±8.38%, Figure 2K). In its active form, SMAD2/3 shuttles into the cell nucleus and activates the transcription of pro-fibrotic genes (as seen in Figure 2C). Overall, this set of results demonstrates that iPSC-cardiac fibroblasts can be obtained from induced pluripotent stem cells and induced to acquire a pro-fibrotic phenotype *in vitro*.

**Figure 2.**
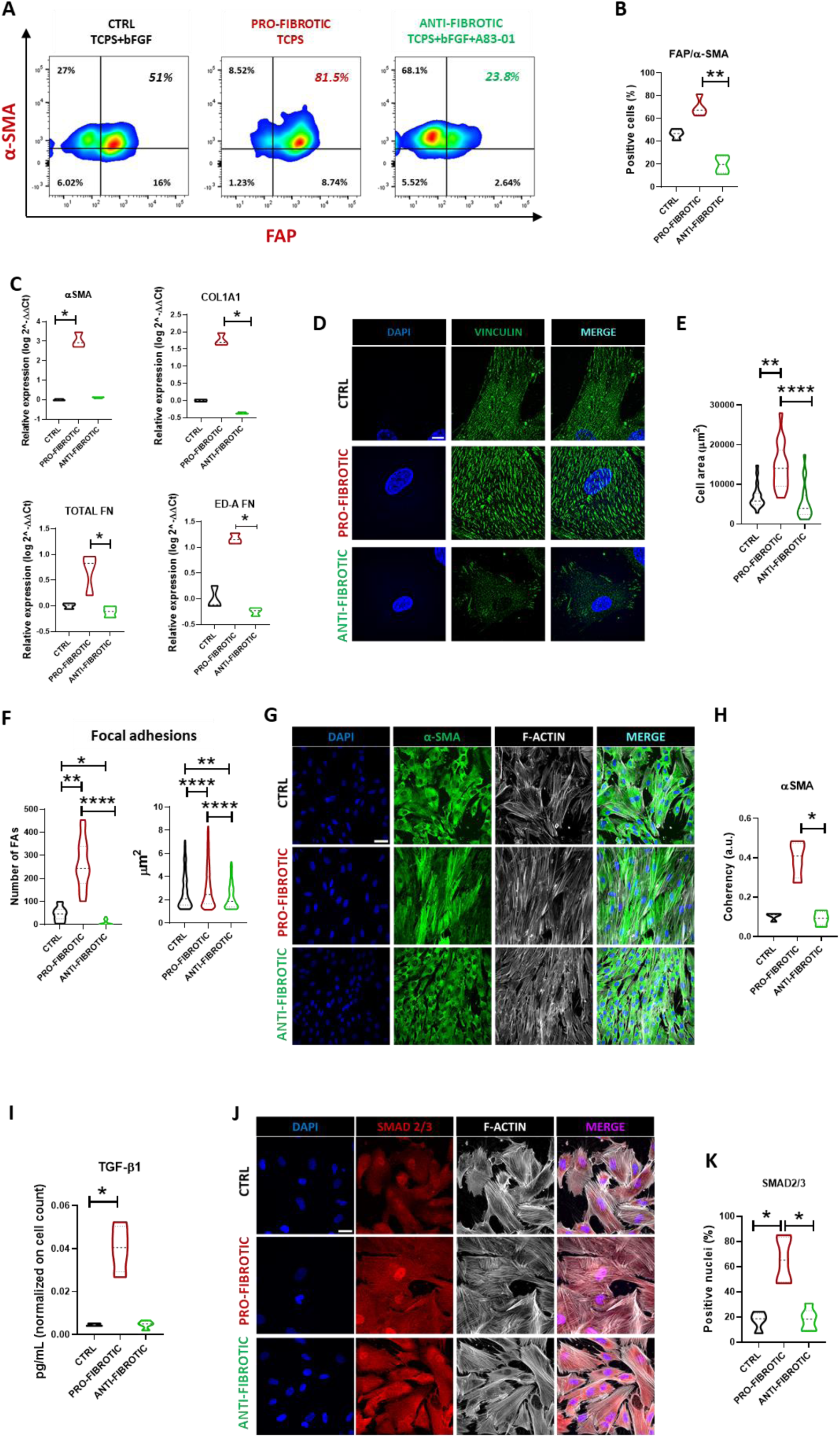
Activation of iPSC-derived cardiac fibroblast to the myofibroblast phenotype. **(A)** Representative contour plot images of the expression of the indicated markers of activated cardiac fibroblasts in ctrl (left), pro- (middle) and anti-fibrotic (right) iPSC-derived cardiac fibroblasts. **(B)** Violin plot representation of the expression of the indicated markers in iPSC-derived cardiac fibroblasts. Values are indicated as percentage of α-SMA/ FAP double positive cells in ctrl, pro- and anti-fibrotic iPSC-derived cardiac fibroblasts (N=3, n=4). **=p < 0.01. Data presented as mean ± standard deviation. **(C)** Violin plot representation of the expression of the indicated markers in iPSC-derived cardiac fibroblasts. Values are indicated as log 2^-ΔΔCt; fold change compared to ctrl (N=3). One-way ANOVA followed by Kruskal-Wallis test, *p<0.05. Data presented as mean ± standard deviation. **(D)** Representative super-resolution microscopy images of focal adhesion protein vinculin expression (green) in ctrl, pro- and anti-fibrotic iPSC-derived cardiac fibroblasts. Nuclei are counterstained with DAPI (blue) (Scale bar: 10µm). **(E)** Violin plot representation of the area of single ctrl, pro- and anti-fibrotic iPSC-derived cardiac fibroblasts. One-way ANOVA followed by Kruskal-Wallis test (N=3; n=17-19). **p<0.01, ****p<0.0001. Data presented as mean ± standard deviation. **(F)** Violin plot representation of focal adhesion (FA) number (left) and area (right) in ctrl, pro- and anti-fibrotic iPSC-derived cardiac fibroblasts. One-way ANOVA followed by Kruskal-Wallis test (N=3; n≥100). *p<0.05, **p<0.01, ****p<0.0001. Data presented as mean ± standard deviation. **(G)** Representative confocal microscopy image of α-SMA (green) and F-actin (grey) in ctrl, pro- and anti-fibrotic iPSC-derived cardiac fibroblasts. Nuclei are counterstained with DAPI (blue) (Scale bar: 50µm). **(H)** Violin plot representation of stress fibres coherency in ctrl, pro- and anti-fibrotic iPSC-derived cardiac fibroblasts as obtained by confocal image analysis. The values are expressed as arbitrary units (a.u.). One-way ANOVA followed by Kruskal-Wallis test (N=3, n=4), *=p<0.05. Data presented as mean ± standard deviation. **(I)** Violin plot representation of TGF-β1 concentration in the supernatant of ctrl, pro- and anti-fibrotic iPSC-derived cardiac fibroblasts. The values are normalized to cell count. One-way ANOVA followed by Kruskal-Wallis test (N=4), *p<0.05. Data presented as mean ± standard deviation. **(J)** Representative confocal images of ctrl, pro- and anti-fibrotic iPSC-derived cardiac fibroblasts stained for SMAD2/3 (red) and F-ACTIN (grey). The nuclei are counterstained with DAPI (blue). (Scale bar: 30µm). **(K)** Violin plot representation of the percentage of ctrl, pro- and anti-fibrotic iPSC-derived cardiac fibroblasts found positive for nuclear SMAD 2/3 in confocal microscopy imaging (N=3, n=4-6). One-way ANOVA followed by Kruskal-Wallis test, *p<0.05. Data presented as mean ± standard deviation.

### Activated iPSC-cFbs deposit fibrotic ECM

ECM remodelling in the heart is orchestrated by cFbs, which drive modifications in its composition, nanostructure and mechanics [^54^]. Therefore, we set at investigating how the phenotype switch induced by iPSC-cFbs activation affected their ability to synthesise and deposit the ECM. For this purpose, iPSC-cFbs were cultured until they reached 100% of confluency (day 7). After a week, cellular components were removed by treating the culture dish with a decellularization solution containing ammonium hydroxide and Triton-X. The dECMs obtained from pro-fibrotic, anti-fibrotic and ctrl iPSC-cFbs were first characterized by scanning electron microscopy (SEM) to study their ultrastructure. SEM analysis revealed that the dECM produced by the fibroblasts exposed to pro-fibrotic stimuli was composed by thick fibres, whereby the ctrl and anti-fibrotic dECMs were characterized by an interconnected web of thin elements (Figure 3A).

**Figure 3.**
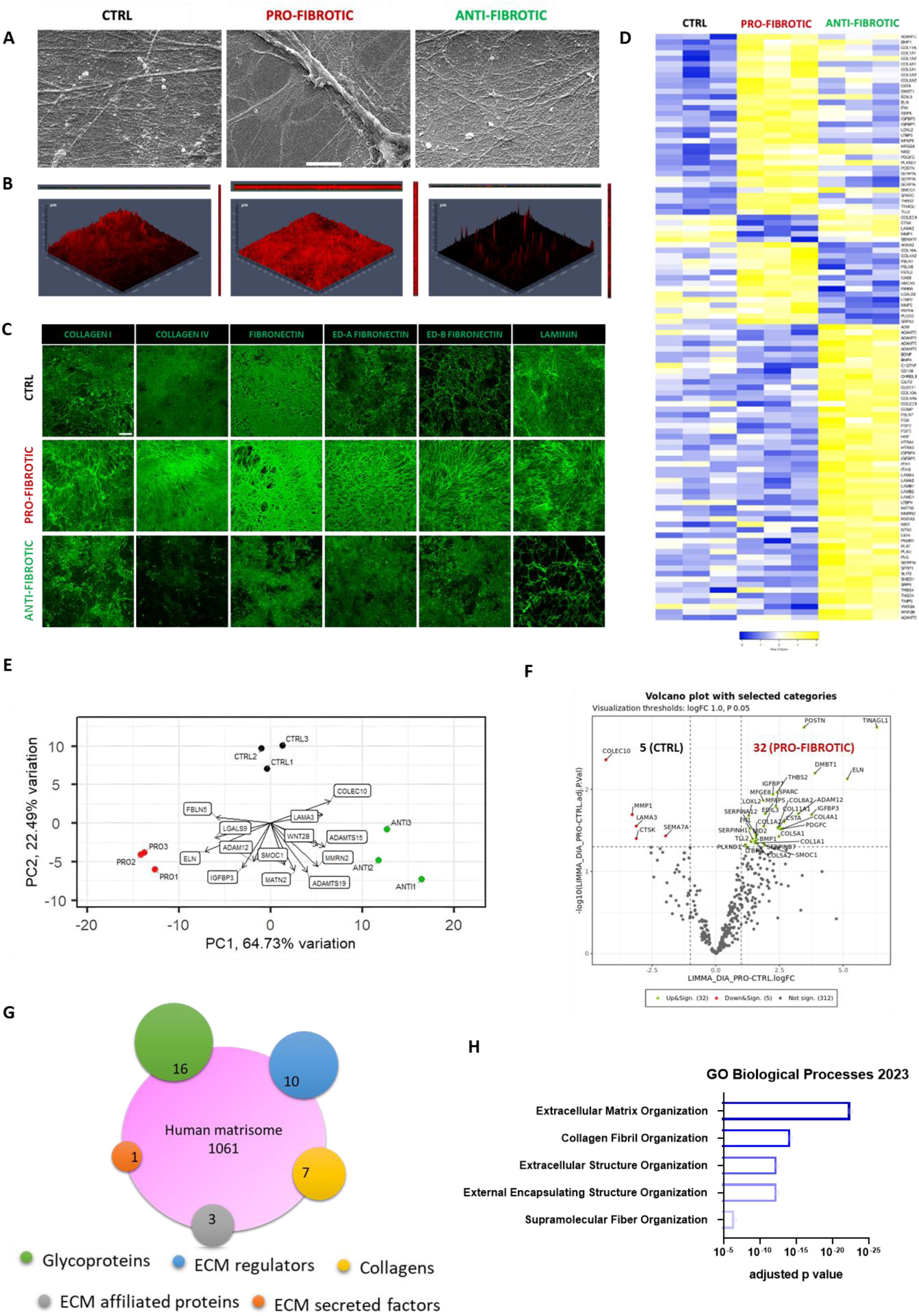
The extracellular matrix deposited by activated cardiac fibroblasts shows a fibrotic signature. **(A)** Representative scanning electron microscopy images showing the architecture of the dECMs deposited by cardiac fibroblasts obtained from iPSCs and exposed to ctrl, pro- and anti-fibrotic culture conditions (Scale bar: 5µm). **(B)** Representative orthogonal view of dECMs derived from ctrl, pro-, and anti-fibrotic iPSC-derived cardiac fibroblasts stained for collagen I (red). The images were obtained by z-stack confocal pictures. **(C)** Representative confocal images of the expression of the indicated proteins (green) in dECMs deposited by ctrl, pro-, and anti-fibrotic iPSC-derived cardiac fibroblasts (Scale bar: 50 µm). **(D)** Heatmap representation of the matrisome proteins found in dECMs obtained from ctrl, pro-, and anti-fibrotic iPSC-derived cardiac fibroblasts and clustered according to their expression pattern. The colour code is adjusted row-by-row based on the row Z-score values (-2 ≤ Z-score ≤ 2). **(E)** Principal component analysis of the clustering of dECMs deposited by ctrl, pro-, and anti-fibrotic iPSC-derived cardiac fibroblasts based on the expression of matrisome proteins. Key components of the matrisome driving the clustering of each of the categories are shown. The length of the arrows weighs the contribution of each protein (N=3). **(F)** Volcano plot representation of the proteins regulated proteins in pro-fibrotic compared to ctrl dECM. The proteins significantly upregulated are shown in green, while those downregulated are indicated in red colour (padj < 0.05, log2Fc > ǀ1ǀ). **(G)** Venn diagram representation of the relative abundance of proteins belonging to the indicated sub-categories of the matrisome found differentially regulated in ctrl vs pro-fibrotic dECM. **(H)** Barplot representation of the indicated Gene Ontology categories (Biological Processes, *GO:0030198; GO:0030199; GO:0043062; GO;0045229*) found significantly regulated in pro-fibrotic vs. ctrl dECM based on the differential expression of matrisome proteins. The graph was obtained from ENRICHR database [^109^] (padj < 0.05, log2Fc > ǀ1ǀ).

The analysis of the three-dimensional structure of the dECMs by confocal microscopy after staining for collagen I demonstrated that while ctrl and anti-fibrotic dECMs appeared to be composed by a thin layer of ECM proteins, fibrotic dECM was – instead – characterized by a dense honeycomb of intertwined fibres (Figure 3B). Of notice, the dECM deposited by iPSC-cFbs grown in anti-fibrotic conditions was unevenly distributed on the well, with entire areas being barely covered by the adhesion proteins (Figure 3B). Next, we characterized the dECM generated by activated, ctrl and anti-fibrotic cFbs by staining them with antibodies directed against ECM structural and basal components (collagen I and IV, fibronectin, ED-A and ED-B fibronectin isoforms, and laminin - Figure 3C). Confocal images indicated an enhanced ECM component deposition in the pro-fibrotic condition as compared to the ctrl and anti-fibrotic counterparts. DAPI counterstaining confirmed that the DNAse treatment following iPSC-cFbs decellularization successfully removed cellular components (data not shown).

Next, we harvested the dECMs and analysed their chemical composition *via* mass spectrometry. PCA indicated that the dECMs obtained from three different biological replicates clustered coherently, thereby indicating that the different treatments generated homogeneous and reproducible preparations (Supplementary Figure 2A). A total of 352 matrisome components were identified by MS in the isolated dECMs (Supplementary Table 1), among which we focused on the ECM proteins differentially regulated in the three conditions. Heatmap representation showed a stark difference in the matrix composition among the three treatments (Figure 3D and Supplementary Figure 2B). Bio-informatics analysis weighing the contribution of given components of the dECMs to the clustering of the samples revealed that proteases like ADAM12, ADAMTS19 and ADAMTS15 drove the formation of discrete and separated clusters of pro-fibrotic and anti-fibrotic dECMs (Figure 3E). These results indicate that ECM remodelling is an integral part of the process leading to the formation of fibrotic ECM.

We next delved into the list of the ECM proteins found by MS and established that they mostly belonged to: i) structural elements, among which glycoproteins (POSTN, FN, ELN, FBLN5, THS4) and collagens (COL1A1, COL1A2, COL11A1) could be found; ii) basal lamina elements (COL4A1, COL4A2, LAMA4, LAMB1, LAMC1); iii) growth factors (proteins from TGFβ, FGF, VEGF, BMP and PDGF families); iv) proteins involved in ECM remodelling such as metalloproteinases (MMP1, MMP2, MMP11, MMP19), metalloproteinases inhibitors (i.e., TIMP1, TIMP2, TIMP3), lysyl oxidase (LOX), and LOX-like proteins (LOXL 1-4) (see Figure 3D). We focused on the matrisome proteins found differentially regulated between the ctrl and pro-fibrotic dECMs and identified the upregulation of 32 proteins in the pro-fibrotic condition compared to the control. In contrast, only 5 ECM components were up-regulated in the dECM produced by iPSC-cFbs in ctrl conditions (Figure 3F). In detail, components belonging to all the matrisome categories were identified including 16 glycoproteins, 10 ECM regulator proteins, 7 collagens, 3 ECM-affiliated proteins, and 1 ECM-secreted factor (Figure 3G and Supplementary Figure 2B). Interestingly, the data indicated that the pro-fibrotic ECM was enriched in proteins such as fibulin 5 (FBLN5, log FC = 2.57), fibronectin (FN, log FC = 1.34) and periostin (POSTN, log FC = 3.46). These proteins play a crucial role in fibrosis development since they represent the most abundant non-collagenous matricellular proteins produced by cFbs acting as ligands for integrin binding and essential for collagen fibrillogenesis [^67,68^]. In addition, the increased expression of collage 4 alpha 1 (COL4A1, log FC = 2.89) and collagen 1 alpha 1 (COL1A1, log FC = 1.54) confirmed the complexity of both the basal lamina and interstitial matrix in the fibrotic cFbs-dECM [^69^]. The increased levels of the collagen-binding osteonectin (SPARC, log FC = 2.36) suggest a further grade of complexity of fibrotic dECM [^70^]. Moreover, elastin (ELN), a protein conferring elasticity to the tissue and known for its involvement in cardiovascular diseases progression, was found strongly up-regulated in fibrotic dECM (log FC = 5.16) [^71^]. On the contrary, MMP1, one of the main effectors of ECM remodelling, was down-regulated in fibrotic cFbs-dECM (log FC = 3.28, ctrl vs fibrotic), suggesting increased dynamics in collagen degradation of the ECM in ctrl [^72^].

To further describe the changes happening in the fibrotic ECM, we examined the GO annotations of the deregulated matrisome components. The results obtained showed that these proteins play a major role in structuring the *ECM architecture* (GO:0030198; GO:0030199; GO:0043062; GO:0045229) by regulating the formation and organization of *supramolecular fibres* (GO:0097435) (Figure 3H). Overall, these data highlight the ability of iPSC-cFbs to produce a rich ECM, which at least partially resembles the complexity and the protein content of the human cardiac ECM. Moreover, pro-fibrotic stimuli promote the positive regulation of components responsible for reinforcing the ECM structure and architecture characteristic of the pathophysiology of heart fibrosis [^32^].

### Pro-fibrotic dECM is characterized by altered mechanical properties

Cardiac extracellular matrix remodelling in consequence of pathological stimuli induces changes in the topography and the mechanical properties of the myocardium. In turn, the changes in the 3D architecture and the altered mechanics of the myocardium affect CM contractility and hamper the organ function [^73^]. We measured the micro-mechanic signature of the dECMs using the high throughput nanoindentation platform Pavone (Optics11Life), which allows for the association of local micro-mechanics with the topographic map of the matrices. The dECMs obtained from iPSC-cFbs cultured under pro-fibrotic, ctrl and anti-fibrotic conditions displayed different microtopography. In particular, the fibrotic dECM showed an irregular surface with ridges that reached the maximum thickness of 13 µm. The surface was characterized by the presence of several peaks, which recalled the presence of large fibres observed *via* confocal and scanning electron microscopy (see Figures 3A, B). On the contrary, the dECM deposited by ctrl cells displayed a smoother surface composed by peaks which reached the maximum thickness of 11 µm. Finally, the surface of the dECM obtained from iPSC-cFbs treated with anti-fibrotic stimuli was flatter and thinner, reaching a maximum height of 5 µm (Figure 4A).

**Figure 4.**
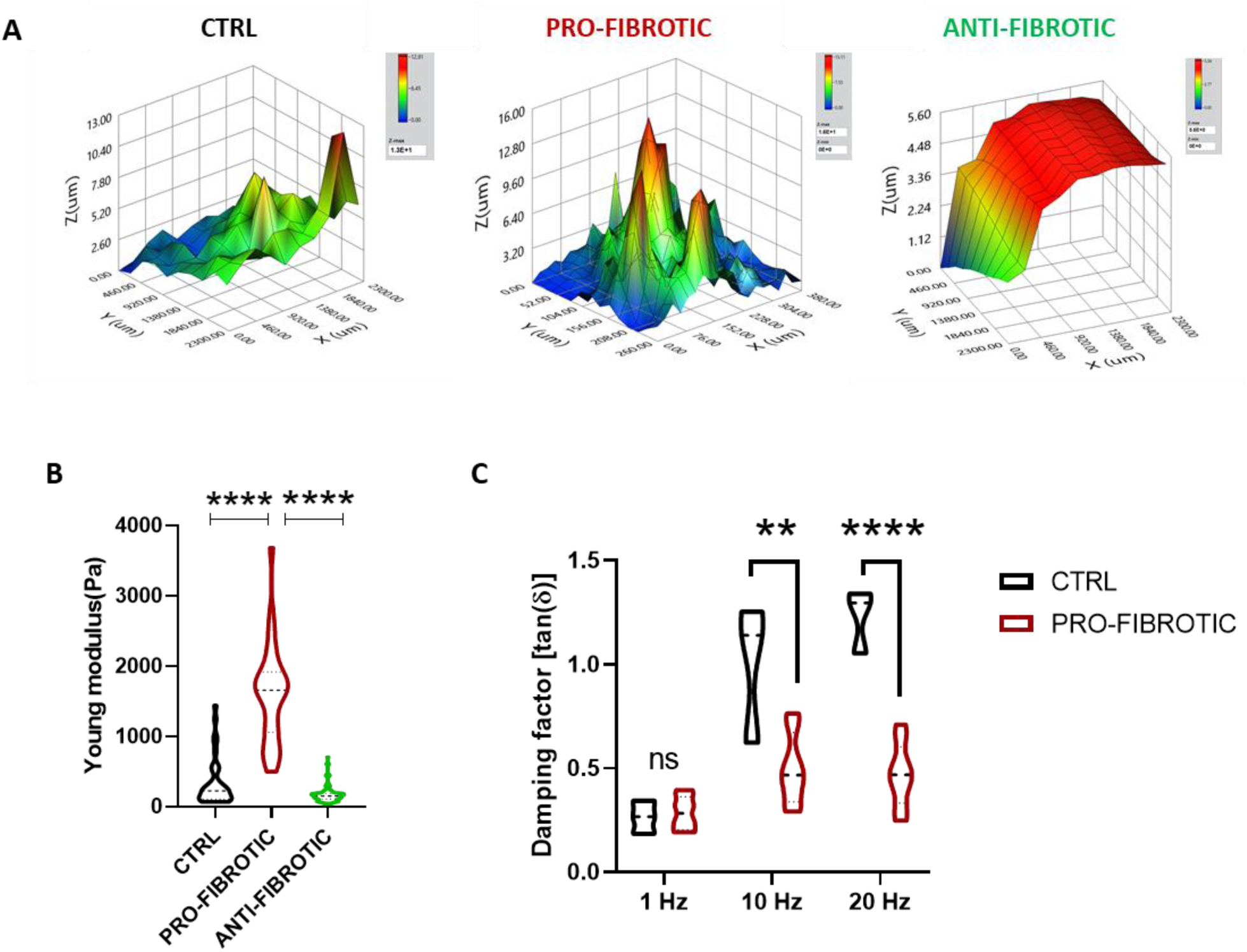
Micromechanics analysis of the dECMs deposited by iPSC-derived cardiac fibroblasts. **(A)** Representative colour-coded maps showing the surface topography of the dECMs deposited by ctrl (left), pro- (middle) and anti-fibrotic (right) iPSC-derived cardiac fibroblasts. The colour code is meant to highlight dECM surface irregularities. **(B)** Violin plot representation of Young’s Modulus values (Pa: Pascal) obtained for ctrl, pro-, and anti-fibrotic dECMs by nanoindentation. (N=3, n≥30). One-way ANOVA followed by Kruskal-Wallis test. ****p<0.0001. Data presented as mean ± standard deviation. **(C)** Violin plot representation of the viscoelastic modulus (Damping factor: tanδ) as obtained after indentation of ctrl and pro-fibrotic dECMs at the indicated frequencies (Hertz: Hz). Multiple t-test validated by Two-stage linear step-up procedure of Benjamini, Krieger and Yekutieli (N=3, n=3) **p<0.01, ****p<0.0001. Data presented as mean ± standard deviation.

The same nano-indentation platform was next used to measure the stiffness and the viscoelasticity of the dECMs. The data obtained showed that all tested samples displayed values of Young’s Modulus in the hundreds of Pascal range, which falls within the range of elasticities reported in previous *in vivo* studies [^32^]. More importantly, the Young’s Modulus of the dECM generated in the pro-fibrotic condition encountered a significant 5-fold increase when compared to that measured in ctrl and a 7-fold upturn as compared to the anti-fibrotic dECM (pro-fibrotic 1593.21 ± 692.32 Pa vs. ctrl 337.40 ± 339.26 Pa vs anti-fibrotic 170.54 ± 110.69 Pa) (Figure 4B). These results are in good agreement with reports coming from *ex vivo* hearts, where stiffening has been described [^32^].

Finally, we performed dynamic measurements to analyse the viscoelastic response of the dECMs obtained from iPSC-cFbs cultured in pro-fibrotic and ctrl conditions following oscillation at different frequencies (1, 10, 20 Hz). Due to the reduced thickness and the uneven distribution of the proteins in the anti-fibrotic dECM (see Figure 4A), viscoelastic measurements could not be performed on that type of sample. This methodology allows to quantify the viscoelastic material response by separately identifying the elastic and viscous components, hence determining the viscous/elastic ratio of its properties expressed as the damping factor tan (δ) [^74^]. Higher tan (δ) values represent higher contributions of viscoelastic components, hence predicting the presence of higher energy dissipating components within a given sample. The analysis demonstrated that the δ factor measured in the pro-fibrotic condition shows meagre, non-significant increases at higher frequencies, suggesting that the elastic component was predominant in this sample compared to the ctrl dECM, which displayed a significant viscoelastic change at increasing frequencies (Figure 4C). Altogether, these results indicate that changes in the mechanical properties of the dECMs could be obtained by exposing iPSC-cFbs to pro-fibrotic conditions. Fibrotic dECM was found to have a higher Young’s Modulus and to be more elastic than the control matrix, which appeared to respond to dynamic stimulations as a more viscous material.

### iPSC-cFbs-derived pro-fibrotic dECMs favour iPSC-CMs survival *in vitro*

Changes in cardiac ECM composition and mechanics are known to impact on postnatal CM phenotype and function [^75, 76,77^]. Similar effects have been shown to occur in pluripotent stem cell-derived CMs *in vitro [*^78^]. To investigate the effects of the dECM deposited by iPSC-cFbs under pro-fibrotic conditions on the phenotype and functionality of contractile CMs, we developed an isogenic *in vitro* model of heart fibrosis entirely relying on iPSCs differentiation. First, we differentiated cFbs from induced pluripotent stem cells, we exposed them to pro-fibrotic or ctrl conditions and isolated the secreted dECMs through the decellularization process described above. Next, we differentiated the same iPSC line into contractile CMs on Matrigel®-coated plates. At day 20, we enzymatically dissociated the beating clusters and re-plated them onto either ctrl or fibrotic dECMs (Supplementary Figure 3A). The purity of the CM population obtained was assessed before transferring them to the dECMs by staining with the CM-specific marker cardiac Troponin T (cTnT). Flow cytometry analysis showed that the protocol yielded a 65.77 ± 5.72% of cells expressing cTnT, confirming CM enrichment in culture (Supplementary Figure 3B).

Next, we monitored the iPSC-CMs on the dECMs. Two days after seeding, brightfield microscopy inspection confirmed that iPSC-CMs cells adhered to both ctrl and fibrotic dECMs and resumed their beating activity (Supplementary Video 1 and Supplementary Video 2). This result was of interest since iPSC-CMs were not able to adhere to uncoated TCPS plate (data not shown). We protracted the cultures until day 30, noticing that the iPSC-CMs did not delaminate from neither the ctrl nor the fibrotic dECM, while they displayed the typical elongated morphology and expressed characteristic sarcomeric proteins like sarcomeric α-actinin (Figure 5A). Moreover, we quantified the percentage of CMs on the dECMs by flow cytometry analysis of cTnT at day 30 of culture. The results showed a slight reduction in the percentage of cTnT-positive cells compared to that quantified before the seeding onto the dECMs. Nonetheless, an enrichment in the number of cTnT-positive CMs in the cells cultured on top of the pro-fibrotic dECM was detected compared to the ctrl dECM (fibrotic: 59.9 ± 7.89 % vs ctrl: 55.64 ± 7.69 %, Figure 5B). Given that our previous experiments demonstrated that in standard Matrigel™ culture conditions iPSC-CMs tend to delaminate with the subsequent reduction of the percentage of cTnT positive cells [^79^], this result suggested that the pro-fibrotic dECM stimulate the adhesion of cardiomyocytes.

**Figure 5.**
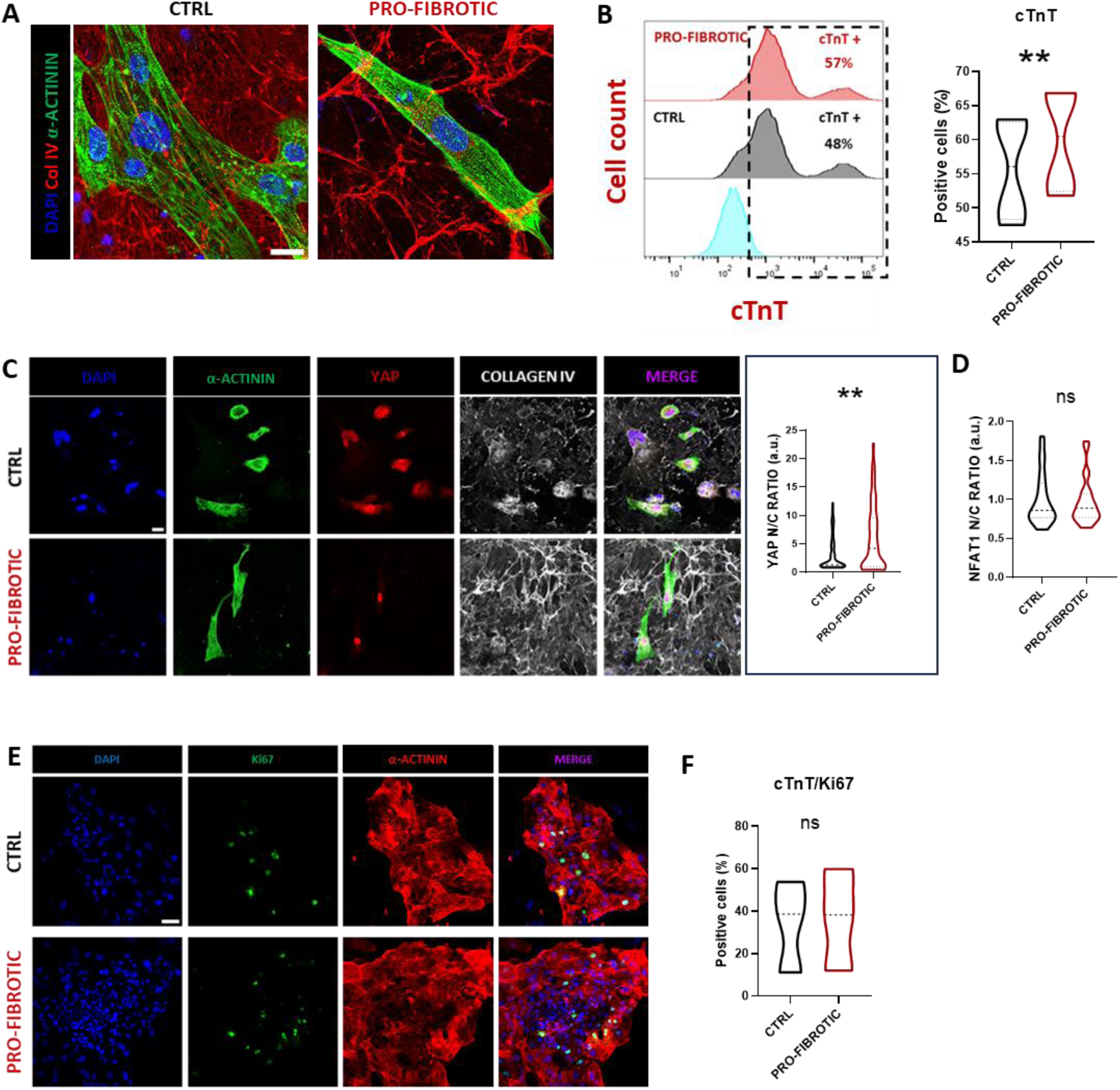
Pro-fibrotic dECM improve iPSC-CMs survival. **(A)** Representative confocal images of iPSC-derived cardiomyocytes cultured for 10 days on ctrl or pro-fibrotic dECMs and stained for α-actinin (green) and collagen IV (Col IV, red). Nuclei were counterstained with DAPI (blue) (scale bar: 20µm). **(B)** Histogram representation (left) and relative violin plot quantification (right) of the percentage of cTnT+ cells found in iPSC-derived cardiomyocytes differentiated for 20 days and cultured for 10 additional days on either ctrl or pro-fibrotic dECM, as obtained by flow cytometry analysis. Ratio paired t-test (N=4). **p<0.01, data presented as mean ± standard deviation. **(C)** Representative confocal images (left) of iPSC-derived cardiomyocytes differentiated for 20 days and cultured for 10 additional days on either ctrl or pro-fibrotic dECM and stained for α-actinin (green), YAP (red) and Col IV (grey). Cell nuclei were counterstained with DAPI (blue) (Scale bar: 30 µm). Violin plot representation (right) of YAP nucleus/cytoplasm ratio in iPSC-derived cardiomyocytes differentiated for 20 days and cultured for 10 additional days on either ctrl or pro-fibrotic dECMs. The values are indicated as arbitrary unit (a.u.). Non-parametric Mann-Whitney test (N=2, n>100). **p<0.01. Data presented as mean ± standard deviation. **(D)** Violin plot representation of NFATC1 nucleus/cytoplasm ratio in iPSC-derived cardiomyocytes differentiated for 20 days and cultured for 10 additional days on either ctrl or pro-fibrotic dECMs. The values are indicated as arbitrary unit (a.u.). Non-parametric Mann-Whitney test (N=2; n=25-28). N.S.=non-significant, data presented as mean ± standard deviation. **(E)** Representative confocal images of iPSC-derived cardiomyocytes differentiated for 20 days and cultured for 10 additional days on either ctrl or pro-fibrotic dECMs and stained for α-actinin (red) and Ki67 (green). Nuclei were counterstained with DAPI (blue). **(F)** Violin plot representation of the co-expression of cTnT and Ki67 in iPSC-derived cardiomyocytes differentiated for 20 days and cultured for 10 additional days on either ctrl or pro-fibrotic dECMs as obtained from flow cytometry analysis. The values are expressed as percentage of double positive cells. Ratio paired t-test (N=3). N.S.= non-significant, data presented as mean ± standard deviation.

To settle this question, we first quantified the nuclear expression of Yes Associated Protein (YAP). The nuclear translocation of the mechanosensitive Hippo downstream effector has been reported in pathological conditions leading to reduced compliance of the heart ECM [^66,32^], likely as a strategy to promote cardiomyocyte survival in the hostile environment [^80^]. This analysis demonstrated that the use of a pro-fibrotic dECM to culture iPSC-CMs determined a significant increase in YAP nuclear/cytoplasmic ratio in the iPSC-CMs (Figure 5C). Since YAP re-expression in adult cardiomyocytes associates with their hypertrophic response, we also investigated the nuclear translocation of NFAT1, an important factor in adult cardiomyocyte pathological hypertrophy [^81,82^] and found no difference between the conditions tested (Figure 5D and Supplementary Figure 3C). YAP nuclear shuttling has also been correlated to the proliferation of iPSC-CMs [^83^]. So, we stained the same cells for Ki67 as a proxy for cell proliferation and detected no significant difference between its expression in iPSC-derived cardiomyocytes cultured either on pro-fibrotic or control dECM (Figure 5E). This result, obtained by immunohistochemistry, was further confirmed by flow cytometry (Figure 5F).

Altogether, these data indicated that both dECMs obtained from iPSC-cFbs exposed to pro-fibrotic or ctrl stimuli represent suitable culture substrates for iPSC-CMs long term culture. The pro-fibrotic dECM favours cardiomyocytes survival – possibly through YAP nuclear translocation - and does not induce any sign of pathological hypertrophy.

### Pro-fibrotic dECM affects iPSC-CMs morphology, sarcomere assembly and function

Next, we set at unveiling the effects of the pro-fibrotic dECM on iPSC-CMs morphology and function. We thus analysed key morphometric parameters of iPSC-CMs in response to the culture on fibrotic or ctrl dECMs at day 30 in culture. The analysis demonstrated that although no differences could be found in iPSC-CMs cellular and nuclear area (Figure 6A, B), cardiomyocytes cultured on fibrotic dECM were significantly less elongated, as demonstrated by their higher circularity (Figure 6C).

**Figure 6.**
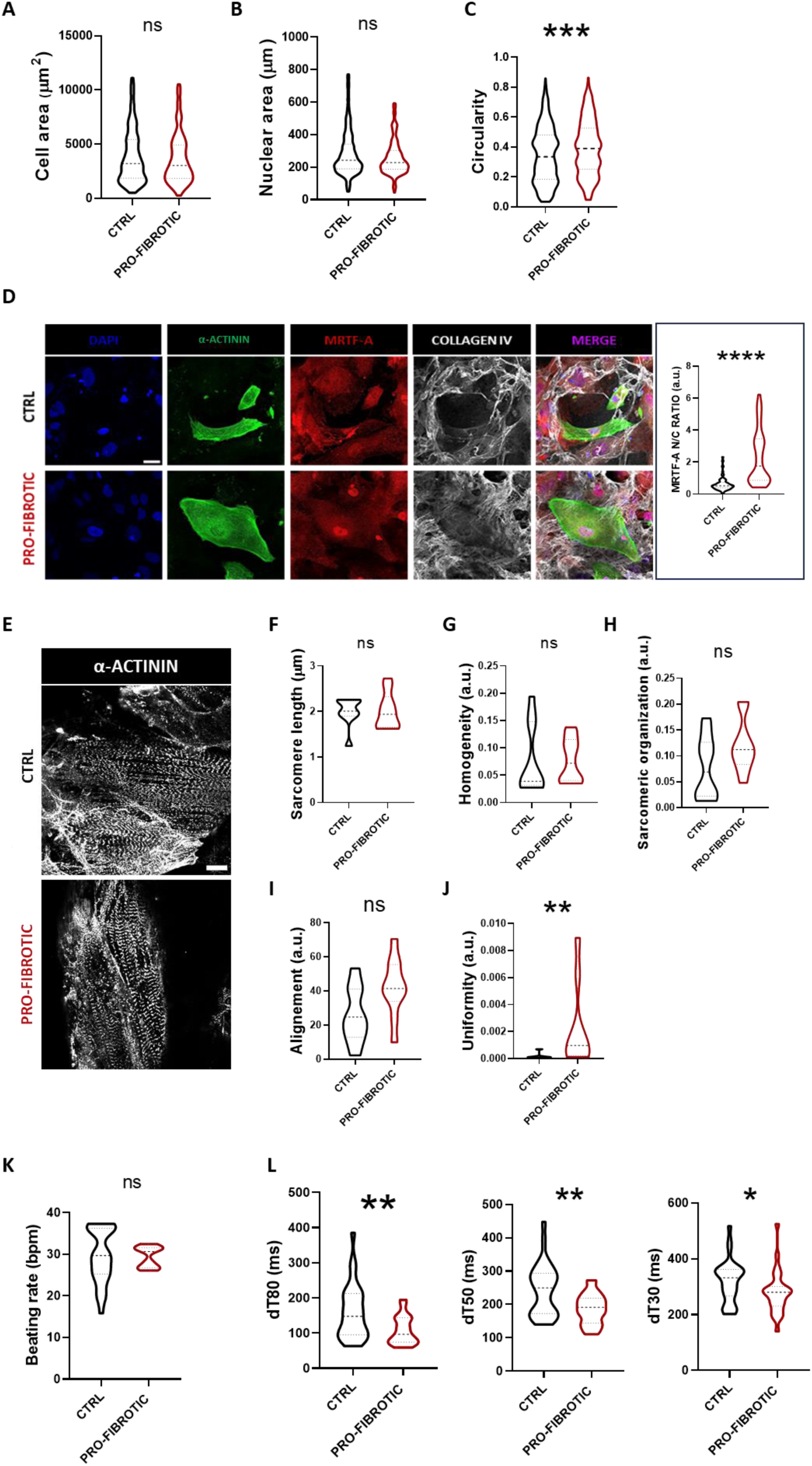
Pro-fibrotic dECM affects iPSC-derived cardiomyocyte sarcomere organization and calcium handling. **(A)** Violin plot representation of the projected area of iPSC-derived cardiomyocytes differentiated for 20 days and cultured for 10 additional days on either ctrl or pro-fibrotic dECMs. **(B)** Violin plot representation of the projected nuclear area of iPSC-derived cardiomyocytes differentiated for 20 days and cultured for 10 additional days on either ctrl or pro-fibrotic dECMs. **(C)** Violin plot representation of the circularity of iPSC-derived cardiomyocytes differentiated for 20 days and cultured for 10 additional days on either ctrl or pro-fibrotic dECMs. Non-parametric Mann-Whitney t-test (N=3; n ≥100). N.S.=non-significant, ***p<0.001. Data presented as mean ± standard deviation. **(D)** Representative confocal images (left) of iPSC-derived cardiomyocytes differentiated for 20 days and cultured for 10 additional days on either ctrl or pro-fibrotic dECMs and decorated with α-actinin (green), MRTF-A (red) and collagen IV (white). The nuclei were counterstained with DAPI (blue). (Scale bar: 30 µm). Violin plot representation of MRTF-A nucleus/cytoplasm ratio in iPSC-derived cardiomyocytes differentiated for 20 days and cultured for 10 additional days on either ctrl or pro-fibrotic dECMs. The values are expressed as arbitrary units (a.u.). Non-parametric Mann-Whitney t-test (N=2; n≥100). ****p<0.0001, data presented as mean ± standard deviation. **(E)** Representative confocal images of iPSC-derived cardiomyocytes differentiated for 20 days, cultured for 10 additional days on either ctrl or pro-fibrotic dECMs and stained for α-actinin (white). Analysis of **(F)** Violin plot representation of the sarcomere length of iPSC-derived cardiomyocytes differentiated for 20 days and cultured for 10 additional days on either ctrl or pro-fibrotic dECMs. **(G)** Violin plot representation of sarcomere homogeneity in iPSC-derived cardiomyocytes differentiated for 20 days and cultured for 10 additional days on either ctrl or pro-fibrotic dECMs. **(H)** Violin plot representation of the sarcomere organization in iPSC-derived cardiomyocytes differentiated for 20 days and cultured for 10 additional days on either ctrl or pro-fibrotic dECMs. **(I)** Violin plot representation of the sarcomere alignment in iPSC-derived cardiomyocytes differentiated for 20 days and cultured for 10 additional days on either ctrl or pro-fibrotic dECMs. **(J)** Violin plot representation of the sarcomere uniformity in iPSC-derived cardiomyocytes differentiated for 20 days and cultured for 10 additional days on either ctrl or pro-fibrotic dECMs. Non-parametric Mann-Whitney t-test (N=3; n=10). N.S.=non-significative; **p<0.01. Data presented as mean ± standard deviation. **(K)** Violin plot representation of the beating rate of iPSC-derived cardiomyocytes differentiated for 20 days and cultured for 10 additional days on either ctrl or pro-fibrotic dECMs. Non-parametric Mann-Whitney t-test (N=3; n=20-30). N.S.=non-significant, data presented as mean ± standard deviation. Data presented as mean ± standard deviation. **(L)** Violin plot representation of calcium decay time (dT80-dT50-dT30) in iPSC-derived cardiomyocytes differentiated for 20 days and cultured for 10 additional days on either ctrl or pro-fibrotic dECMs. Non-parametric Mann-Whitney t-test (N=3; n=27-30). N.S.=non-significant; *p<0.05, **p<0.01. Data presented as mean ± standard deviation.

Recent studies suggested that pluripotent stem cell-derived CMs aspect ratio correlates with the maturation of the sarcomere structures and their ability to generate force [^78^]. Hence, we set at investigating the architecture of the sarcomere assembled by iPSC-CMs grown on the dECMs. Under pathological stress, the integrity of the sarcomeres is guaranteed by myocardin-related transcription factors (MRTFs) nuclear presence [^84^]. The staining of MRTF-A in iPSC-CMs cultured on dECMs and subsequent analysis by confocal microscopy revealed a significant increase in its nuclear/cytoplasmatic ratio in CMs cultured onto pro-fibrotic dECM compared to the ctrl counterpart (Figure 6D).

Next, we adopted an automatic tool for the quantification of key parameters describing sarcomere length and organization in iPSC-CMs grown on either ctrl or fibrotic dECMs. For this purpose, the cells were stained with an antibody against α-actinin to image z-disc spacing in confocal microscopy (Figure 6E). Although no differences in sarcomere length and homogeneity existed between iPSC-CMs cultured on ctrl and fibrotic dECMs (Figure 6F, 6G), the analysis demonstrated a tendency to increase for parameters like sarcomere organization and alignment in pro-fibrotic conditions (Figure 6H, 6I). More intriguingly, the culture on fibrotic dECM induced a significant increase in sarcomere uniformity compared to cells grown onto ctrl dECM (Figure 6J). Altogether, these results indicate that iPSC-CMs responded to stimuli arising from the dECM obtained from activated iPSC-cFbs by increasing their sarcomere alignment and organization, two parameters which are key to their contractile activity.

Lastly, we asked whether the structural changes found in iPSC-CMs cultured onto fibrotic iPSC-cFbs-derived dECMs would also result into alterations in the calcium handling properties in these cells. Although the beating rate of the CMs remained unaffected in response to fibrotic dECMs compared to the ctrl one (fibrotic: 27.80 ± 1.92 bpm vs. ctrl: 29.63

± 3.47 bpm) (Figure 6K), the calcium-binding fluorescent dye Fluo-4 showed that the CMs seeded on pro-fibrotic dECM had a significantly shorter calcium decay time (dt80, dt50, dt30) (Figure 5L). This result indicated that, when cultured on fibrotic dECMs, iPSC-CMs displayed a faster relaxation compared to those cultured on ctrl dECM. No significant difference was - instead - detected in calcium uptake (Supplementary Figure 4A).

In conclusion, these results indicate that the pro-fibrotic dECM alters iPSC-CMs relaxation by regulating their calcium homeostasis [^85^].

## Discussion

Myocardial fibrosis highlights the progression of a multitude of cardiovascular diseases. Its onset and evolution are orchestrated by activated cardiac fibroblasts as an attempt to preserve the integrity of the organ by preventing heart wall rupture. Eventually, this process leads to the deposition of a non-compliant, fibrotic scar tissue which might hamper the organ pump function by impairing CMs contractile activity or the electrical conductance [^1^]. The biomechanical stress associated with pathological ECM remodelling also modifies the transcriptional and post-transcriptional landscape of cardiac cells [^33^].

Unfortunately, no effective treatment exists at present to prevent cardiac fibroblast activation or reverse the detrimental effects of excessive ECM deposition. Current strategies to reduce the impact of cardiac fibrosis include angiotensin (AT)-converting enzyme and AT receptor antagonist, β-blockers, as well as statins and endothelin antagonists. The aldosterone pathway inhibitor Eplerenone has also been adopted to suppress fibrosis onset, while modulators of TGFβ / Smad3 signalling in fibroblast activation are currently under evaluation [^86^].

Among the main limitations on the way to treat myocardial fibrosis is the lack of physiological *in vitro* models able to recapitulate its main characteristics while capturing patient specificity [^87^]. Considering the 3Rs principle of replacement, reduction, and refinement which encourage the progressive substitution of animal experimentation, the use of animal models appears outdated. In fact, besides the ethical and practical concerns associated with such practices [^88, 89,90^], the housing and handling costs associated with them cannot be overlooked. This is also valid for the recent attempts at generating *in vitro* platforms that rely on the use of decellularized tissues and organs from animals.

The most recent attempts at modelling cardiac fibrosis rely on microfluidic platforms combining biochemical and mechanical stimuli to induce cardiac fibroblasts activation to myofibroblasts [^91^], or multicellular systems based on the co-culture of cardiomyocytes and cardiac fibroblasts of various origin [^92, 93^].

Here we established and validated a protocol to obtain activated cardiac fibroblasts from human induced pluripotent stem cells and devised a convenient strategy to harvest intact fibrotic decellularized matrices. When checked against the composition of human pathological heart ECM, these matrices show remnants of a cardiac fingerprint [^32^].

Finally, we used the fibrotic matrices for long time culture of iPSC-derived cardiomyocytes. By culturing CMs on dECMs generated from the same iPSC line, we established an isogenic cardiac fibrosis model suited for patient-specific pre-clinical studies.

We first demonstrated that iPSCs can be induced to differentiate into fibroblast-like cells that express high percentages of cardiac (GATA4, TBX5) and fibroblast-specific markers (CD90, FSP and DDR2) [^20, 94, 95, 96, 97^]. Similar to what was previously shown in post-natal cardiac fibroblasts [^62^], these cells became spontaneously activated to the myofibroblast phenotype when cultured on a stiff substrate. The acquisition of the contractile phenotype could be confirmed by the increased cell area, the formation of α−SMA-rich stress fibres in concomitance with the expression of FAP. Moreover, this result was corroborated by detecting the accumulation of FAs, the nuclear shuttling of SMAD2/3 effector and the upregulation of fibrotic genes like COL1A1, FN and its splicing variant ED-A in cells cultured on stiff substrate.

Since iPSC-derived cardiac fibroblasts cultured on TCPS released in the supernatant a fair amount of TGF-β1 - the growth factor known to be the primary driver of cardiac fibroblasts activation *in vivo - [*^98, 99, 55, 73^], we used a pharmacological inhibitor of TGF-β receptor-family (A83-01) to test how it affects iPSC-cFbs activation, alone or in combination with bFGF [^35, 100^]. We showed that both the experimental conditions were effective in reducing iPSC-derived cardiac fibroblasts activation, as shown by reduced cell area and the disappearance of the characteristic α−SMA-rich stress fibres. These alterations were significantly more pronounced when the dual treatment with bFGF and the TGF-β1 inhibitor was applied.

The abnormal deposition of ECM proteins by cFbs is the main hallmark of fibrotic condition [^5,6^]. Therefore, we established a decellularization procedure suited to remove all cellular components from the iPSC-cFbs culture, while preserving the intact dECM for further studies [^101^]. Interestingly, the dECM from pro-fibrotic cFbs was enriched in interstitial fibres, strongly represented by fibronectin pathological splicing isoforms, organized in an intricated honeycomb of crosslinked filaments. On the contrary, the treatment with TGF-β1 inhibitor in combination with bFGF hampered the ability of cFbs to secrete abundant ECM and to crosslink it, so that its organized and harmonic structure was mostly lost. The changes observed in dECM architecture resulted from differences in protein composition which we validated using mass spectrometry. This analysis showed an upregulation in the dECM produced by fibroblasts grown onto the stiff substrate of structural ECM proteins such as COL1 and COL4, FBLN5, POSTN and glycoproteins involved in the critical inward transduction of extracellular cues like FN. Interestingly, for the same experimental conditions we found proteins involved in the determination of ECM elasticity like SPARC and ELN [^8, 102, 103^] to be significantly upregulated.

We next adopted a high throughput strategy to sample by nanoindentation the micromechanical properties of the dECMs produced by iPSC-cFbs cultured onto TCPS and treated or not with bFGF alone or in combination with TGF-β receptor-family inhibitor. While confirming that iPSC-cFbs deposited a thick dECM characterized by a multitude of peaks at the microscale, the analysis revealed that its stiffness lied in the kPa range, which is consistent with the physiological Young’s Modulus values found for the myocardium [^104^]. It is important to remark that the values we report here (1-2 kPa) might be slightly lower than the values found *in vivo* (15-40 kPa) since they are obtained in the absence of the confounding element of the contractile cells.

Interestingly, the treatment with bFGF alone or in the presence of A83-01 significantly caused a 5- to 7-fold reduction in the stiffness of the dECM. This effect was due to a reduction in the complexity of the surface of the dECM. We also performed dynamic measurements that indicated a more pronounced viscous response for the dECMs deposited by cells treated with bFGF compared to those grown in the absence of inhibition. Although we were not able to obtain any data from the dECMs produced by cFbs exposed to bFGF in concomitance with A83-01, the results here reported could serve as a playground to predict the viscoelastic behaviour of the anisotropic myocardium ECM, which has not been fully estimated so far. In fact, few studies on the viscoelastic properties of the heart to date were performed on intact swine ventricular myocardial tissue, which was found to exhibit viscoelastic properties. While the main elastic contribution in the heart is thought to be coming from titin functioning as a spring, the viscous input might be ascribed to the collagen component of the ECM [^105, 106^].

ECM stiffening has been described as a key event in the fibrotic process and decisive to drive the impairment of cardiomyocyte contractile activity in the long run [^77, 78^]. Therefore, we tested the possibility of using the dECMs obtained from cells cultured on TCPS (pro-fibrotic) or the same cells treated with bFGF (ctrl) as substrates for the long-term culture of iPSC-derived cardiomyocytes. Remarkably, we demonstrated that it is possible to culture iPSC-CMs - usually grown on Matrigel™ or Cultrex™ derived from rodent tumours lacking cardiac-specific cues [^24,25,23,26^] - onto fibrotic and ctrl matrices secreted by iPSC-cFbs for up to 30 days without signs of delamination of the beating cells.

Instead, by stimulating the cFbs to acquire the myofibroblast phenotype and by culturing cardiomyocytes obtained from the same iPSC line on top of dECM secreted by the cFbs, we produced an isogenic cardiac fibrosis *in vitro* model. By using this strategy, we showed that the pro-fibrotic dECM triggers a mechanosensing response in cardiomyocytes derived from iPSCs, fosters their survival, and impacts on their morphology and sarcomere assembly. More importantly, the use of fibrotic dECM as culture substrate led to modified cardiomyocyte relaxation rate by affecting their calcium homeostasis. Changes in heart relaxation are typical of the diastolic dysfunction, with recent reports suggesting that myocardial fibrosis contributes to its pathogenesis [^107^]. This process is characterized by abnormal ventricular distensibility, relaxation and filling, and it precedes heart failure [^108^].

## Conclusion

In conclusion, by using decellularized matrices obtained from iPSC-derived cardiac fibroblasts induced to acquire a pro-fibrotic phenotype, we generated a versatile *in vitro* tool to study the impact of pathological ECM remodelling and drug experimentation on cardiomyocyte function in a patient-specific setting. This approach might be in future used to overcome the limitations of current culture substrates.

## Acknowledgements and Fundings

This project has received funding from the European Union’s Horizon 2020 research and innovation programme under the Marie Skłodowska-Curie grant agreement No 860715, by the King’s BHF Centre for Excellence Award (RE/18/2/34213) and by Marie Curie H2020-MSCA-IF-2020 MSCA-IF-GF “MecHA-Nano”, Grant Agreement No 101031744. CIISB, Instruct-CZ Centre of Instruct-ERIC EU consortium, funded by MEYS CR infrastructure project LM2023042, is gratefully acknowledged for the financial support of the LC-MS/MS measurements at the CEITEC Proteomics Core Facility. Computational resources were provided by the e-INFRA CZ project (ID:90254), supported by MEYS CR. We are thankful to Helena Ďuríková for helping with cell culture, Romana Vlčková and Hana Dulová for the continuous technical support. We are grateful to CELLIM facility (CEITEC) for their contribution in acquiring cell images with super resolution microscopy. Cartoons were created by BioRender (https://www.biorender.com/).

## Authors’ contributions

F.N., S.F., M.C. and G.F. conceptualized the study and drafted the manuscript. F.N. designed, performed the experiments and analysed the data. S.F., M.C., J.O.D.L.C., M.A., D.P.S., V.V., S.P., G.P., E.S., D.R., A.S.M., M.R. assisted in designing, carrying out and interpreting experiments. Z.Z., D.P., and V.P. performed dECM by mass-spectrometry and analysed the data. L.B., and M.B. performed micro-mechanics measurements and analysed the raw data.

## Declaration of interest

Luca Bersanini and Malin Becker are employees of Optics11 Life B.V

## Supplementary Figures

**Supplementary Figure 1.**
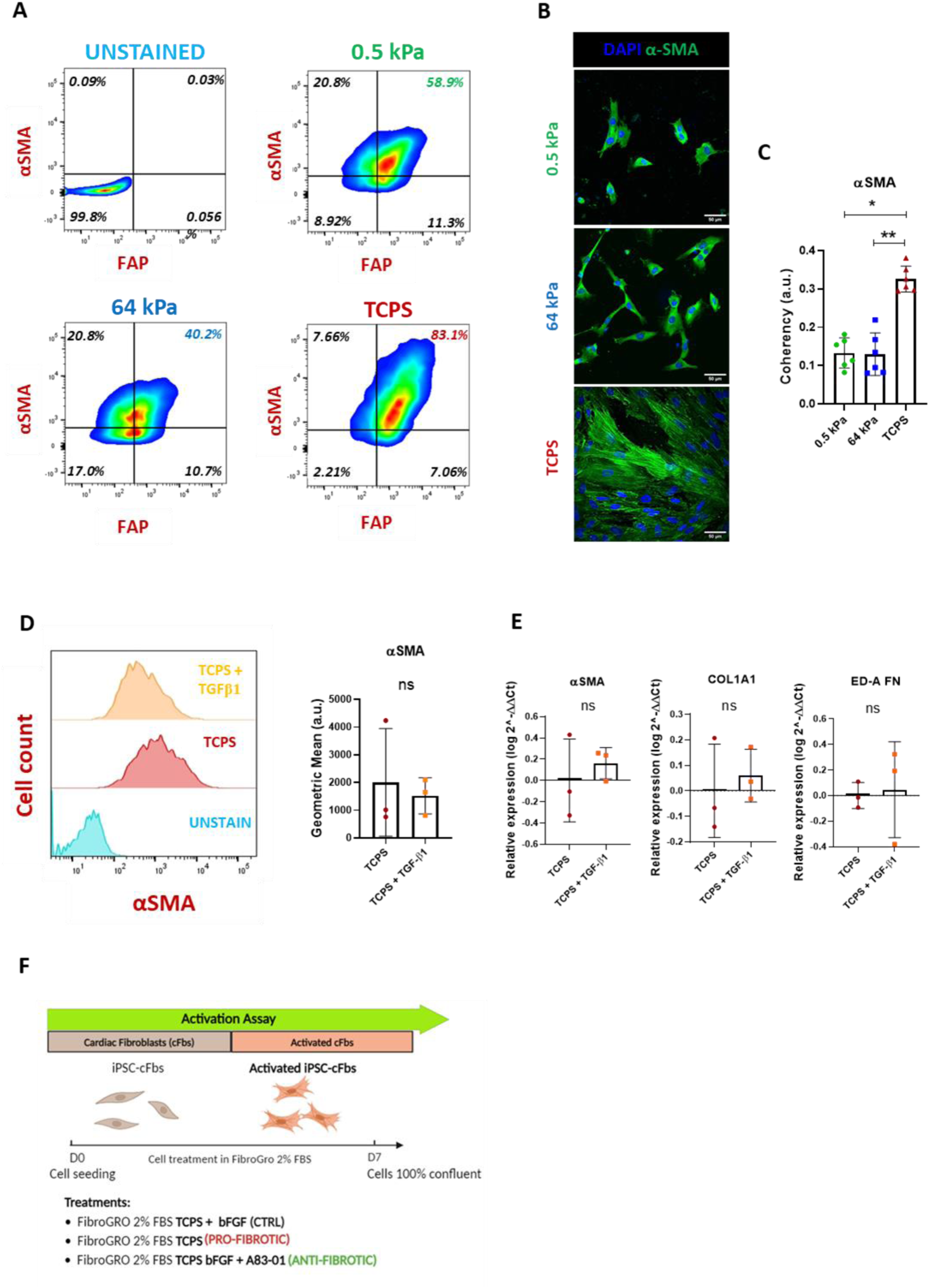
Effect of substrate on iPSC-derived cardiac fibroblast activation. **(A)** Representative contour plots obtained by flow cytometry and describing the expression of Fibroblast Activated Protein (FAP) and alpha smooth muscle actin (α-SMA) in iPSC-derived cardiac fibroblasts cultured for 7 days on 0.5 kPa and 64 kPa substrates or on tissue culture polystyrene (TCPS). **(B)** Representative confocal images of iPSC-derived cardiac fibroblasts cultured for 7 days on 0.5 kPa and 64 kPa substrates or on tissue culture polystyrene (TCPS). The cells were stained for α-SMA (green) and the nuclei were counterstained with DAPI (blue) (scale bar: 50 µm). **(C)** Barplot representation of α-SMA fibres alignment coherency in iPSC-derived cardiac fibroblasts cultured for 7 days on 0.5 kPa and 64 kPa substrates or on tissue culture polystyrene (TCPS). The values are expressed as arbitrary units (a.u.). One-way ANOVA followed by Kruskal-Wallis test (N=3, n = 6). *p<0.05, **p<0.01, data presented as mean ± standard deviation. **(D)** Representative histogram plot (left) and its relative barplot representation (right) of α-SMA expression in iPSC-derived cardiac fibroblasts cultured for 7 days on tissue culture polystyrene (TCPS) alone or in the presence of TGFβ-1 as obtained by flow cytometry. The curves in the histogram plot indicate cell counts, while α-SMA values in the barplot are expressed as intensity geometric mean. One-way ANOVA followed by Kruskal-Wallis test (N=3). **(E)** Barplot representation of the expression of the indicated genes in iPSC-derived cardiac fibroblasts cultured for 7 days on tissue culture polystyrene (TCPS) alone or in the presence of TGFβ-1 as obtained by RT-qPCR. The results are expressed as log 2^-(ΔΔCt) normalized to TCPS. One-way ANOVA followed by Kruskal-Wallis test (N=3). N.S.= non-significant. **(F)** Schematic representation of the iPSC-derived cardiac fibroblast activation procedure adopted in this study.

**Supplementary Figure 2.**
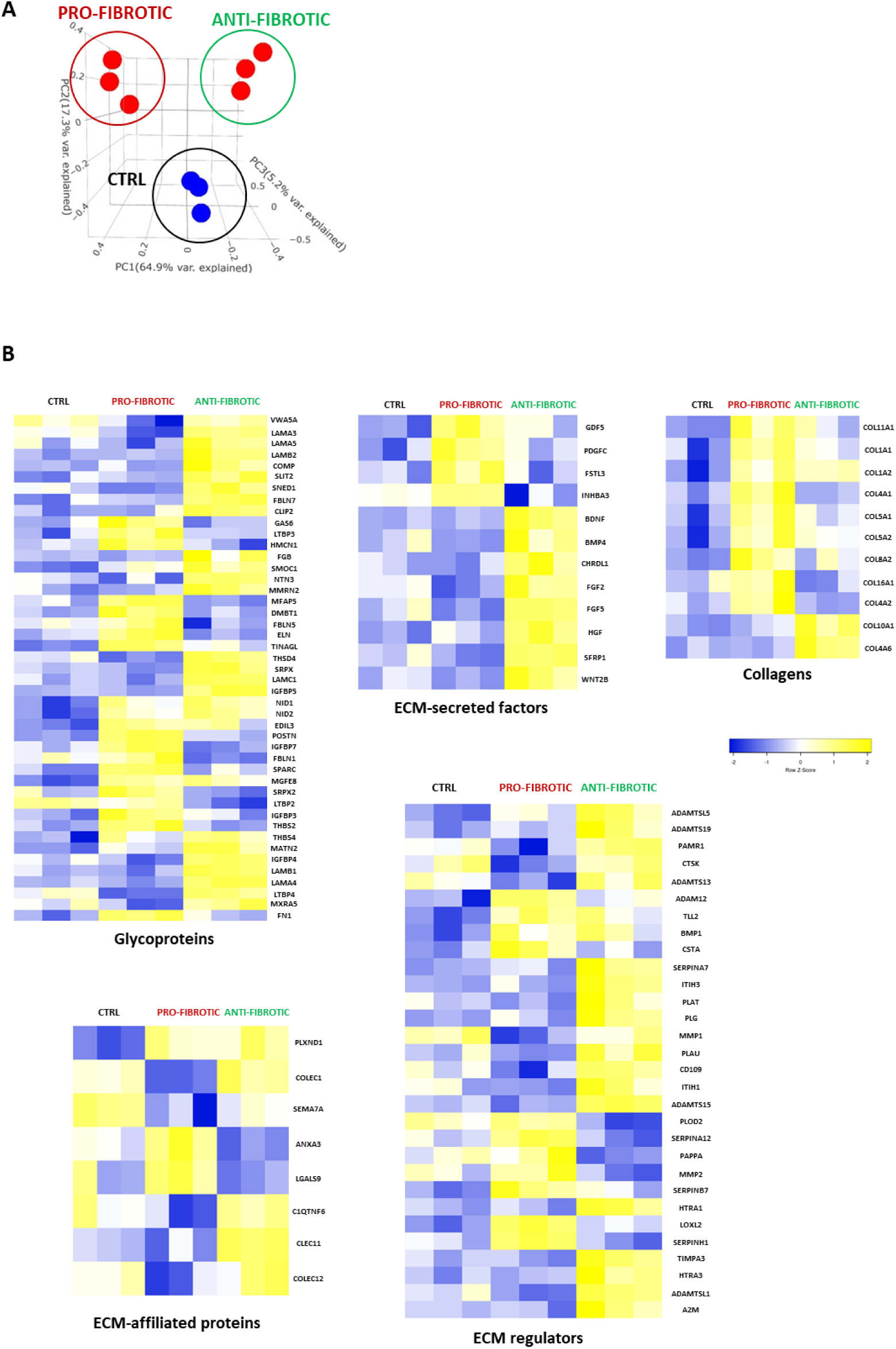
Mass spectrometry characterization of the dECMs obtained from iPSC-derived cardiac fibroblasts. **(A)** 3D principal component analysis (PCA) of the protein composition of dECMs deposited by ctrl, pro- and anti-fibrotic iPSC-derived cardiac fibroblasts, as obtained by Mass Spectrometry. The analysis was performed *via* BIOJUPIES [^110^]. **(B)** Heatmap representation of the indicated genes belonging to the annotated categories (glycoproteins, ECM regulators, ECM-affiliated proteins, ECM-secreted factors) found expressed in dECMs deposited by ctrl, anti-fibrotic and pro-fibrotic iPSC-derived cardiac fibroblasts.

**Supplementary Figure 3.**
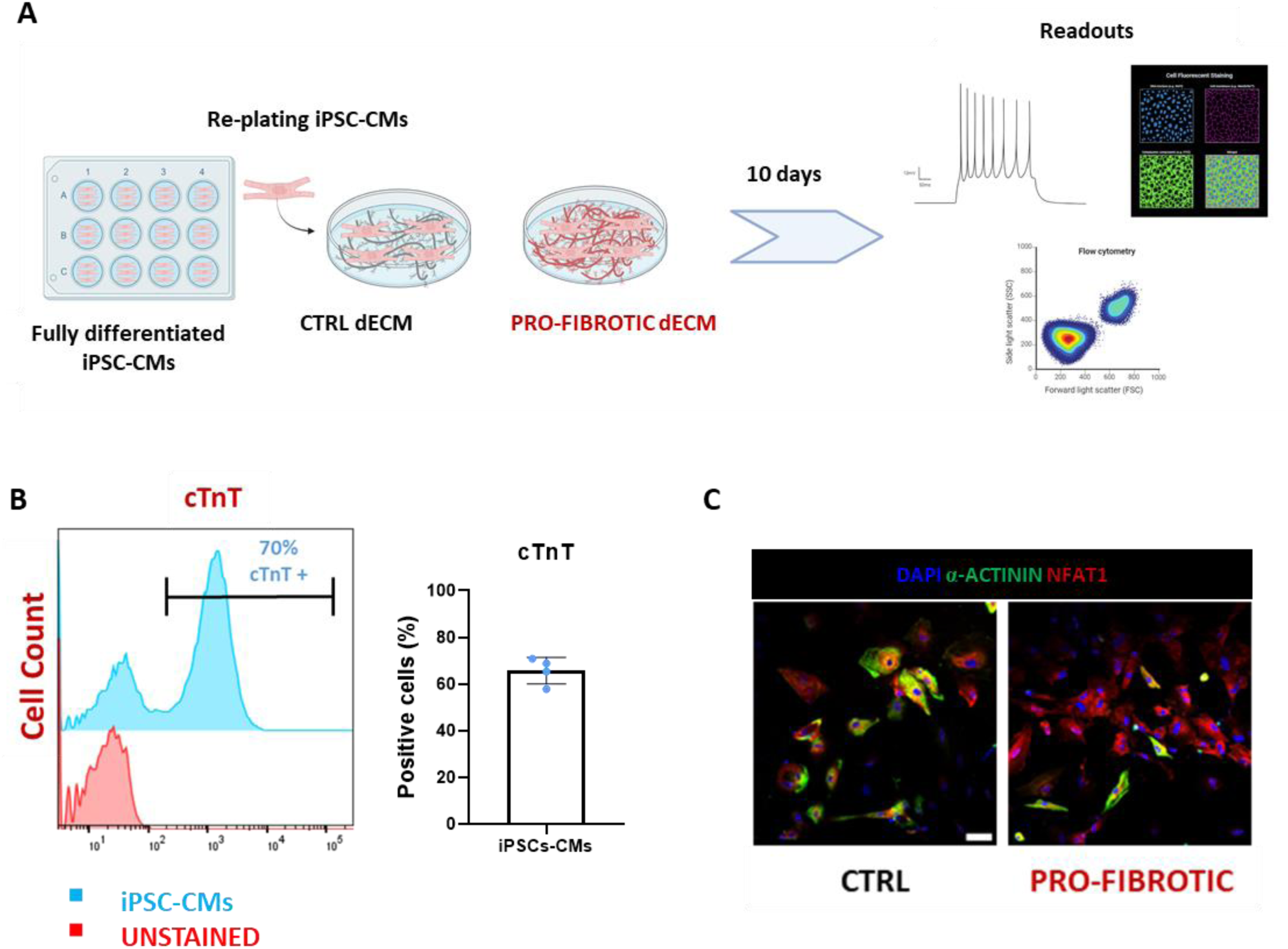
Characterization of iPSC-derived cardiomyocytes cultured onto the dECMs deposited by iPSC-derived cardiac fibroblasts. **(A)** Schematic representation of the protocol adopted in the study. **(B)** Representative histogram plot (left) and relative barplot quantification (right) of the percentage of cells expressing cTnT in iPSC-derived cardiomyocytes at day 20 of differentiation. The data were obtained by flow cytometry for 4 independent experiments. Data presented as mean ± standard deviation. **(C)** Representative confocal images of iPSC-derived cardiomyocytes cultured on ctrl and pro-fibrotic dECM and stained for α-actinin (green) and NFAT1 (red). The nuclei were counterstained with DAPI (blue) (Scale bar: 50 µm).

**Supplementary Figure 4.**
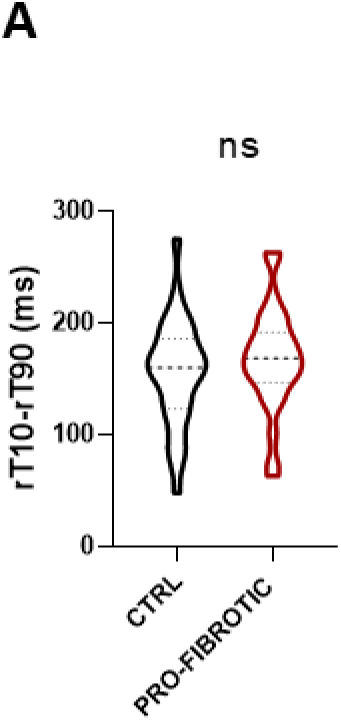
Calcium uptake in iPSC-derived cardiomyocytes cultured onto dECMs obtained from iPSC-derived cardiac fibroblasts. **(A)** Violin plot representation of the calcium uptake time (rT90-rT10) in iPSC-derived cardiomyocytes cultured onto dECMs deposited from iPSC-derived cardiac fibroblasts exposed or not to pro-fibrotic stimuli. Non-parametric Mann-Whitney test. (N=3; n = 10). N.S.=non-significant. Data presented as mean ± standard deviation.

